# α-Arrestins maintain phospholipid balance and Atg18 distribution to permit efficient autophagy

**DOI:** 10.1101/2022.08.03.502596

**Authors:** Ray W. Bowman, Sydnie Davis, Eric M. Jordahl, Karandeep Chera, Nejla Ozbaki-Yagan, Jonathan Franks, Sarah Hawbaker, Annette Chiang, Omer Acar, Donna Beer-Stoltz, Allyson F. O’Donnell

## Abstract

Cells selectively reorganize their membrane proteome in response to stressors via selective protein trafficking. The α-arrestins, a family of conserved protein trafficking adaptors, bind to select membrane proteins and interact with the ubiquitin ligase Rsp5. The α-arrestins recruit Rsp5 to its membrane protein substrates, permitting their ubiquitination and endocytosis. To identify new α-arrestin functions, we performed a genetic screen to isolate mutants that alter α-arrestin-mediated resistance to rapamycin, a drug that inhibits TORC1. Interestingly, loss of many of the *ATG* genes, which encode the machinery needed for the self-degradative process of autophagy, disrupted α-arrestins’ ability to promote growth on rapamycin. Herein we define a genetic network linking α-arrestins to autophagy. We show autophagy impairment in the absence of select α-arrestins, with increased autophagosome lifetimes and delayed/reduced delivery of autophagosomes to the vacuole. The α-arrestin mutants that impeded autophagy had vacuole morphology defects and increased vacuolar retention of Atg18, a member of the PROPPIN family that is needed to maintain vacuole shape and facilitate lipid transfer to expanding autophagosomes. Atg18 binds phosphatidylinositol 3 phosphate (PI3P) and phosphatidylinositol 3,5-bisphosphate (PI(3,5)P_2_), and we observed increased PI3P on the vacuole membrane in α-arrestin mutants. The levels of Vps34 and Fab1, the kinases responsible for the generation of PI3P and PI(3,5)P_2_, respectively, were also elevated at vacuole membranes in cells lacking α-arrestins. We posit that altered phospholipids in the vacuolar membrane form the basis for the Atg18-Atg2 mislocalization and autophagy defect. These data demonstrate a previously unappreciated link between the α-arrestins and autophagy, expanding the functional impact of these trafficking adaptors in responding to nutrient stress.

**Author Summary:** Cells survive nutrient starvation by degrading parts of themselves through the process of autophagy. During autophagy, cells make a double membrane, known as an <u>a</u>uto<u>p</u>hagosome (AP), around bits of cytoplasm or organelles. The AP and its engulfed material are delivered to the vacuole, an organelle that helps break down proteins and lipids. These materials can then be used as building blocks to generate the essential components needed for the cell to survive starvation. For cells to undergo efficient autophagy, they need α-arrestins, a group of proteins important for deciding where membrane proteins localize. In cells lacking α-arrestins, the AP forms slowly, likely due to a problem in growing the AP membrane. This results in less material being delivered to the vacuole via APs when cells do not have α-arrestins. This study defines a new role for α-arrestins in promoting AP formation and starvation survival.

## Introduction

Cells require the appropriate nutrient balance to grow and thrive. In nutrient-replete conditions, the Target of Rapamycin Complex 1 (TORC1) is a kinase that helps drive constructive metabolic processes, allowing for cell growth and division [1]. Key to TORC1 regulation of the metabolic program is its ability to stimulate synthesis of macromolecules, but equally important is TORC1’s inhibition of destructive metabolic processes, like autophagy or the endocytic turnover of nutrient transporters [2–9]. In times of nutrient starvation, TORC1’s repressive functions are lost and macro-autophagy (hereafter referred to as autophagy), the process whereby cells non-selectively engulf portions of the cytosol or organelles in a double-membrane structure referred to as an <u>a</u>uto<u>p</u>hagosome (AP), is activated. APs are formed using a dedicated machinery encoded by the <u>a</u>utophagy-related <u>g</u>ene family (*ATG*) [10, 11]. APs deliver cellular material to the vacuole (the yeast equivalent of the lysosome), via fusion with the vacuolar membrane. In parallel, TORC1-mediated inhibition of endocytosis is lost under starvation conditions and many membrane proteins at the <u>p</u>lasma <u>m</u>embrane (PM), including key nutrient transporters, are endocytosed and trafficked to the vacuole for degradation [12]. In both starvation-induced autophagy and endocytosis, the components delivered to the vacuole are broken down to generate building blocks that fuel essential processes during nutrient depletion [13, 14]. There is a dynamic interplay between starvation-induced endocytosis and autophagy; productive endocytosis and breakdown of transporters in the vacuole is required to maintain the cell during early phases of starvation, prior to full autophagic induction [12]. While a global defect in endocytosis impedes autophagy progression, we do not yet understand how more selective defects in membrane protein endocytosis impact autophagy progression.

Nutrient- and TORC1-regulated endocytosis of membrane transporters relies heavily on the α-arrestins, a class of selective protein trafficking adaptors conserved from yeast to man [15–18]. In *Saccharomyces cerevisiae*, where α-arrestin function is best described, nutrient deprivation or rapamycin treatment inactivates TORC1, alleviating the phospho-inhibition on α-arrestins — in part through the activation of the downstream protein phosphatase Sit4 — and increasing transcription of some α-arrestins [2,18–20]. Dephosphorylated α-arrestins stimulate starvation-induced endocytosis of many types of membrane proteins, including amino acid permeases [17,18,21–26]. To achieve membrane protein endocytosis, the α-arrestins selectively bind to membrane proteins, and while the interface between α-arrestins and their targets has yet to be fully mapped, acidic patches in the cytosolic portions of membrane proteins are important for this recognition [27–31]. α-Arrestins also interact with the ubiquitin (Ub) ligase, Rsp5, serving as a molecular bridge between the ligase and membrane cargo [15,17,32–35]. When α-arrestins are bound to membrane proteins, Rsp5 can ubiquitinate both the substrate and the α-arrestin (Fig 1A) [15,17,18,36,37].

**Fig 1.**
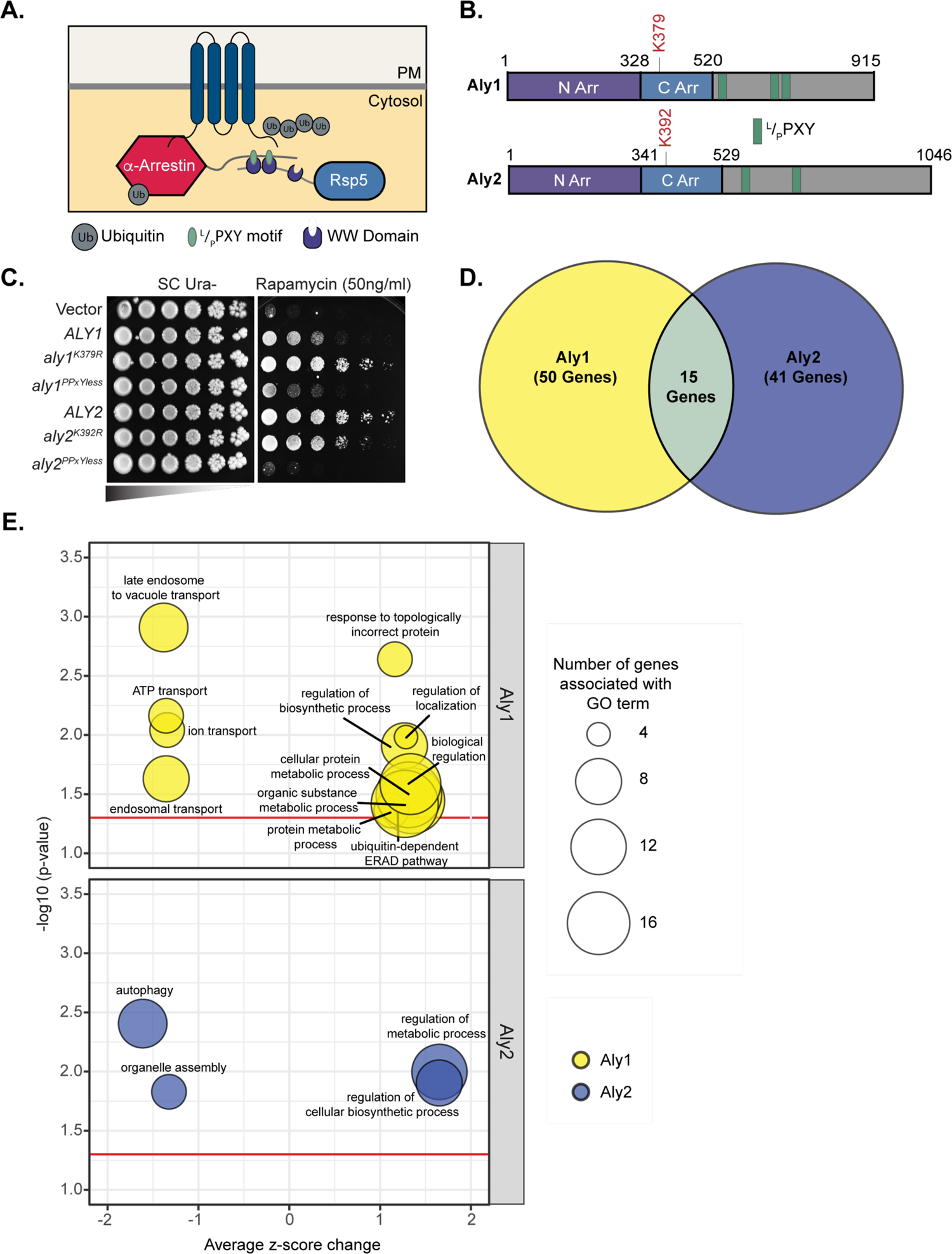
Genetic screen of the ScUbI library identifies modifiers of α-arrestin-mediated resistance to rapamycin. (**A**) Model of α-arrestin function. α-Arrestins bind selectively to membrane proteins and interact with the Rsp5 ubiquitin ligase. α-Arrestin ^L^/_P_PXY motifs (green) interact with the WW domains of Rsp5 (purple). Rsp5 can ubiquitinate (Ub in grey) the α-arrestin, which regulates α-arrestin function, or the membrane protein, which helps stimulate its endocytosis. (B) A schematic of α-arrestins Aly1 and Aly2 depicting the N- and C-terminal arrestin-fold domains (purple and blue, respectively), ^L^/_P_PXY motifs (green) and sites of mono-ubiquitination [1]. Numbers indicate amino acid positions in coding sequence. (C) Serial dilution growth assays of WT cells containing the pRS426-derived plasmid expressing either nothing (vector) or the indicated α-arrestin on SC medium lacking uracil, with and without 50ng/mL rapamycin, are shown. (D) Venn-diagrams indicating the number of ScUbI library gene deletions that altered Aly1 or Aly2-mediated resistance to rapamycin. (E) Representative gene ontology terms for the genes with the strongest Z-score changes in *ALY1* (top) or *ALY2* (bottom) overexpression backgrounds. The x-axis shows the average Z-score changes for the genes associated with a particular GO term and the y-axis represents the significance of the association. For GO terms with similarities, we only show the parent terms. Circle sizes represent the number of terms associated with the GO terms. Many genes with strong negative Z-score changes in Aly2 are related with autophagy.

Membrane protein ubiquitination serves as a trigger for endocytosis as the ubiquitinated target protein interacts with early-arriving endocytic patch proteins containing ubiquitin-interaction motifs [38–41]. α-Arrestin mono-ubiquitination is thought to be activating, helping the α-arrestin to bind to Rsp5 and assisting in α-arrestin recruitment to membranes [22, 42]. Together, α-arrestin-regulated endocytosis helps control cellular nutrient supply and ensures that, under starvation conditions, the membrane proteome is remodeled to aid in cellular adaptation.

We sought to uncover novel aspects of α-arrestins’ function in response to starvation. We used rapamycin-induced, TORC1 inhibition as a basis for a targeted genetic screen, focusing on genes important for ubiquitination or ubiquitin-related modifiers, including the ubiquitin-like modifiers that regulate autophagy [43, 44]. We found that when we over-expressed two paralogous α-arrestins, Aly1 or Aly2, we induced resistance to rapamycin, suggesting that these factors somehow prevent rapamycin-induced inhibition of TORC1. We were surprised to find that loss of genes required for autophagy often altered the Aly-induced rapamycin resistance. We further examined the genetic interactions between α-arrestins and autophagy components and found strong ties between phagophore membrane nucleation and expansion proteins, including proteins in the <u>p</u>hosphatidylinositol <u>3</u>-<u>k</u>inase (PI3K) complex, which synthesizes the needed <u>p</u>hosphatidylinositol <u>3</u>-<u>p</u>hosphate (PI3P) for incorporation into the growing phagophore, and the phosphoinositide-binding, PROPPIN family member, Atg18, which regulates phosphoinositide balance in the vacuolar membrane and helps in lipid transfer to the growing AP at the <u>p</u>hagophore <u>a</u>ssembly <u>s</u>ite (PAS) [45–52].

Defects in either Vps34 or Atg18 impair autophagic flux [45,46,53–55]. Recent studies demonstrate that mutations that hyperactivate Vps34, thereby increasing cellular PI3P, lead to prolonged AP lifetimes and delayed AP fusion to the vacuole [54]. PI3P is partially responsible for recruiting Atg18, and its partner Atg2, to the PAS [46, 47]. In turn, the Atg2-Atg18 complex along with Atg9, a transmembrane protein localized to cytosolic vesicles that are thought to help form the isolation membrane during autophagy, aid in the supply of lipids to the expanding phagophore membrane and tether the PAS on the vacuole to the <u>e</u>ndoplasmic <u>r</u>eticulum (ER) [49,50,56,57]. The cellular roles of Vps34 and Atg18 expand beyond autophagy, each able to influence retromer-dependent retrograde trafficking, <u>e</u>ndosomal <u>s</u>orting <u>c</u>omplexes <u>r</u>equired for transport (ESCRT) function in vacuole-mediated degradation, and vacuole morphology and fusion events [52,58–63].

Based on our genetic screen, we explored the connection between α-arrestins and autophagy. We found that in the absence of select α-arrestins, including Aly1, Aly2 and Art1 (aka Art6, Art3 and Ldb19, respectively) autophagic flux to the vacuole was delayed [64–66] We further showed by electron microscopy that APs formed in response to rapamycin were smaller in the absence of α-arrestins and there were fewer <u>a</u>utophagic <u>b</u>odies (AB) present in the vacuoles of α-arrestin mutant cells than WT cells. Like the increased AP lifetime observed in hyperactive Vps34 cells [54], in the absence of α-arrestins AP lifetimes were also increased. We found elevated PI3P on the limiting membrane of the vacuole and an enrichment of phosphoinositide binding proteins, Atg2 and Atg18, on vacuoles in cells with disrupted α-arrestin function. Taken together, we propose a model in which α-arrestins are needed to maintain PI3P, and possibly PI(3,5)P_2_, balance on the vacuole membrane. In the absence of select α-arrestins, elevated PIPs allow for aberrant accumulation of Atg18 and Atg2 on the limiting membrane of the vacuole, possibly disrupting the lipid transfer function of Atg18-Atg2 at the PAS to give rise to prolonged AP lifetimes and delays in AP expansion observed in α-arrestin mutants. This work raises new questions about α-arrestin function in cells, highlighting their impact on phosphoinositide distributions, and sets the stage for future studies focused on the metabolic ramifications of disrupting α-arrestin-mediated trafficking and their role in autophagy.

## Results

### Identifying genes that alter Aly-mediated growth on rapamycin

TORC1 is a key signaling regulator, promoting growth in the presence of robust nutrient supply, and preventing catabolic processes [1]. However, nutrient starvation or treatment with the drug rapamycin inhibits TORC1, resulting in a signaling transition that blocks protein synthesis, induces the self-degradative process of autophagy, and inhibits cell growth [1]. Through their role in controlling nutrient transporter trafficking the α-arrestins are key regulators of nutrient homeostasis and several α-arrestins are regulated by TORC1 [2,18,22]. We found that over-expression of α-arrestins Aly1 or Aly2 conferred resistance to rapamycin on solid media (Fig 1C) [32]. In addition to Alys, over-expression of α-arrestins *ART1* or *ART5* also conferred rapamycin resistance, which we explore further below (S2A Fig). The α-arrestins utilize ^L^/_P_PXY motifs, typically located in C-terminal tail extensions after the arrestin-fold domains, to bind the WW-domains of Rsp5 (Fig 1A-B). Aly-induced rapamycin resistance was dependent on their interaction with Rsp5 since PPXYless mutants, where their ^L^/_P_PXY motifs had been converted to PPXG to ablate Rsp5 binding [32], failed to promote rapamycin resistance (Fig 1C). This suggests that α-arrestin-induced trafficking may be important for this activity. α-Arrestins are mono-ubiquitinated, which often activates their endocytic function, or poly-ubiquitinated, which triggers their degradation (Fig 1B) [15,17,18,37,67–70]. We mapped the sites of mono-ubiquitination on α-arrestins Aly1 and Aly2 using a mass spectroscopy approach [71] and generated mutants replacing the lysine residue targeted for mono-ubiquitination with arginine (K379R and K392R in Aly1 and Aly2, respectively). These mutations prevent Aly mono-ubiquitination [71]. Despite being paralogs, the loss of mono-ubiquitination influenced Aly proteins differently, improving or impairing the rapamycin resistance conferred by Aly1 and Aly2, respectively (Fig 1C).

To help define α-arrestin regulators, we next conducted a targeted genetic screen of ubiquitin-related regulatory factors. We constructed the *<u>S</u>accharomyces <u>c</u>erevisiae* <u>Ub</u>iquitin Interactome (ScUbI) library, an array of 323 unique deletion mutants that were each annotated in the *<u>S</u>accharomyces cerevisiae* <u>G</u>enome <u>D</u>atabase (SGD) as being linked to ubiquitin or ubiquitination (*see Materials and Methods*). To help define α-arrestins’ role in nutrient regulation, we leveraged the rapamycin resistance phenotype associated with *ALY* over-expression and screened the ScUbI library for modifiers. As part of an undergraduate laboratory course (see *Materials and Methods*), we compared the growth on rapamycin of yeast over-expressing *ALY1* or *ALY2* to the vector control, and identified a total of 76 gene deletions that significantly altered colony growth in an Aly-dependent manner on rapamycin-containing medium in comparison to vector-expressing cells (ΔV +/- 1.0, see S1 Datafile). 50 and 41 gene deletions altered Aly1- or Aly2-dependent growth, respectively, with 15 of these impacting both α-arrestins (Fig 1D; see S1 Datafile). When we assessed Gene Ontology term enrichment for candidates in our screen, we found that loss of ER-associated degradation (ERAD) genes enhanced α-arrestin-mediated resistance to rapamycin, while loss of autophagy-related genes interfered with the α-arrestin-associated resistance to rapamycin (Fig 1E).

### A network of genetic interactions between autophagy facilitators and α-arrestins

We performed a secondary screen to better define the autophagy-related genes that impact α-arrestin-mediated growth on rapamycin. To broadly assess the genetic links between α-arrestins and autophagy, which utilizes a diverse and dedicated set of machinery belonging to the *ATG* gene family [10, 11], we analyzed gene deletion mutants from different stages of the autophagy pathway. From these studies, we confirmed many of the genetic interactions observed in our ScUbI library screen (including *atg2Δ*, *atg17Δ*, and *atg18Δ*) and found several additional *ATG*-family deletion mutants that altered Aly-mediated rapamycin resistance (including *atg6Δ*, *atg31Δ*, and *atg34Δ)* (Fig 2A-C and S1 Fig). To enable a quantitative comparison of our growth assay results, an example of which is presented for *atg2*Δ in Fig 2A, we designed a Cell Profiler-based analysis pipeline to measure the pixel intensities associated with cell growth in images from these assays (see *Materials and Methods*). The quantified data are presented as a heat map in Fig 2B, where white represents maximum growth and black represents no growth (Fig 2B, and primary data in S1 Fig). For easier interpretation, we then overlay these genetic interactions on a model of the autophagy pathway (Fig 2C).

**Fig 2.**
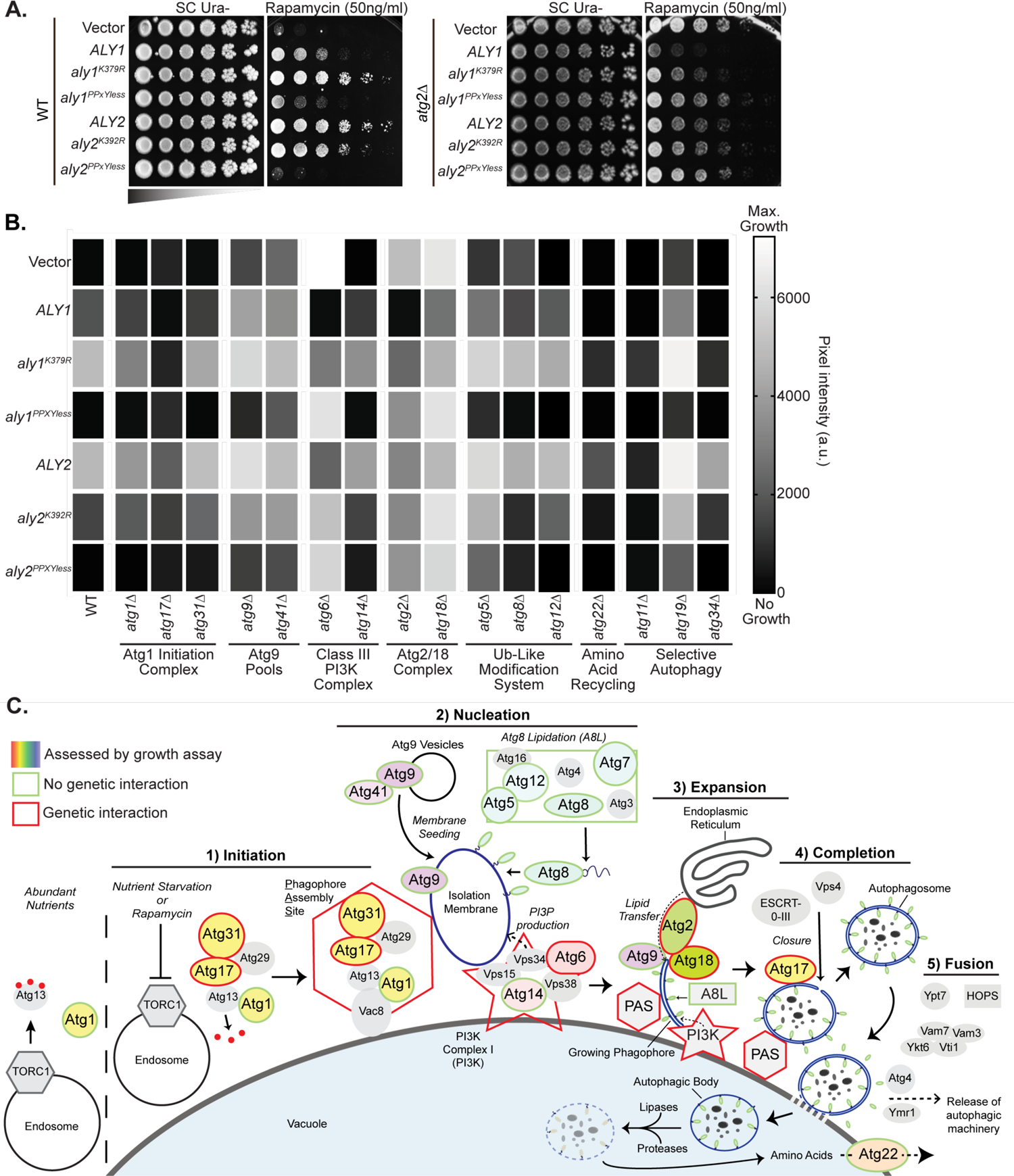
Genetic interactions between α-arrestins and the *ATG* gene family. (A) Serial dilution growth assays of WT and *atg2Δ* cells containing a pRS426-derived plasmid expressing either nothing (vector) or an α-arrestin on SC medium lacking uracil with or without 50ng/mL rapamycin are shown. One representative example of the larger serial dilution growth assay set, which is found in S1 Fig, is provided. (B) Heat map representing quantified (see *Materials and Methods*) serial dilution growth assays (for primary data see S1 Fig) of WT or *atg*Δ cells containing a pRS426-derived plasmid expressing either nothing (vector) or the indicated α-arrestin on SC medium lacking uracil with or without 50ng/mL rapamycin. (C) A simplified diagram of the machinery involved in macroautophagy divided into 5 stages: 1) Initiation, 2) Nucleation, 3) Expansion, 4) Completion, and 5) Fusion. In short, under abundant nutrient conditions, the endosomal population of the TORC1 signaling complex inhibits autophagy through phosphorylation of Atg13 (red circles). Once endosomal TORC1 is inhibited, either by nutrient starvation or treatment with the drug rapamycin, Atg13 is dephosphorylated, allowing for Atg1 activation and the recruitment of other autophagic machinery to form the PAS and initiate macroautophagy (1). Vesicles containing Atg9 are then recruited to the PAS, providing the seed for isolation membrane formation. The autophagy-specific PI3K is also recruited to the PAS, along with machinery to conjugate Atg8 to PE for incorporation into the isolation membrane (2). PI3K activity at the PAS produces PI3P on the isolation membrane, recruiting the Atg18-Atg2 complex. Atg2, bound to Atg9, tethers the tip of the growing phagophore to the ER and facilitates ER-to-phagophore lipid transfer which, along with Atg8 lipidation, drives phagophore expansion (3). Once fully expanded to encapsulate its cytosolic cargo, the phagophore is sealed through activity of Atg17, Vps4, and the ESCRT machinery to form a completed AP (4). The autophagic machinery is then released and Atg8 is cleaved from its PE anchor on the outer autophagic membrane. APs then fuse to the vacuolar membrane through the action of Rab GTPase Ypt7, the HOPS tethering complex, and various SNARE proteins, delivering the inner-membrane-bound autophagic body to the vacuolar lumen (5). There, vacuolar lipases and proteases degrade the autophagic body and its contents, producing free amino acids that are effluxed to the cytosol via Atg22. Shapes filled with color were assessed with the serial dilution growth assays found in (B) and S1 Fig, while those filled with grey were not. Those deletion mutants found to have a genetic interaction with α-arrestins are marked with a red outline, while those that marked with a green outline had no genetic interaction with α-arrestins.

We first evaluated genes involved in the initiation of autophagy. Atg17 is a scaffolding protein for the PAS, acting as a regulatory subunit along with Atg13 to stimulate Atg1 kinase activity and playing a role AP fusion to the vacuole [72, 73]. *ATG17* loss prevented *ALY1* over-expression from conferring rapamycin resistance (Fig 2B, column 3). The loss of *ATG17* also dampened the rapamycin resistance conferred by other α-arrestins (*ALY2*, *ALY2^K392R^, ART5*, and *ART1*) (Fig 2B, S2B-C Fig). Loss of *ATG31*, which binds Atg17 and is similarly needed for autophagy initiation [74–76], also reduced rapamycin resistance for Aly1, Art5 and Art1, showing largely the same trends as *atg17Δ* cells (Fig 2B, column 4 and S2B-C Fig). However, for Aly2, higher rapamycin resistance was observed in the *atg31*Δ cells compared to WT cells (Fig 2B, column 4 and S2B-C Fig). Despite the impact of Atg17 and Atg31 on α-arrestin-induced rapamycin resistance, loss of the *ATG1* kinase, which is essential for autophagy initiation, did not dramatically change the rapamycin-resistance phenotypes (Fig 2B, column 2 and Fig 2C).

We next evaluated genes encoding components found in PI3K Complexes I and II, which facilitate autophagy and endomembrane trafficking, respectively [45,77,78]. Atg6 (aka Vps30) is a subunit of both PI3K Complexes I and II and is required for autophagy [45]. We found that *atg6*Δ cells were resistance to rapamycin, yet, surprisingly, the over-expression of Aly1, Aly2, Art1 or Art5 severely reduced rapamycin resistance in this genetic background (Fig 2B, column 7, Fig 2C, and S2B-C Fig). The Aly K-to-R mutants that block mono-ubiquitination of these α-arrestins behaved similarly to their cognate WT alleles, while the PPXYless mutants, which fail to bind Rsp5, had no impact on the *atg6*Δ-induced resistance to rapamycin. The PI3K Complex I is specific to autophagy and it contains the autophagy specific components, Atg38 and Atg14, instead of Vps38, which is found in PI3K Complex II [45, 79]. Atg14 anchors Atg6 to the PI3K complex and PAS, where the complex produces PI3P on the phagophore’s isolation membrane and recruits the lipid transfer complex, Atg2 and Atg18 [78, 80]. Atg14 recruitment to the PAS relies on Atg13, Atg17, and Atg9 [80]. Unlike Atg6, which is needed for both PI3K Complexes I and II, loss of *ATG14* did not increase rapamycin resistance. However, α-arrestin-mediated rapamycin resistance was dampened in *atg14Δ* cells (Fig 2B, column 8 and Fig 2C).

Downstream of the PI3K complex, Atg2, an autophagy-specific protein of the Vps13 family, functions as a lipid-transfer protein, anchoring the ER to the PAS to permit expansion of the phagophore [51,81,82]. Like *atg6Δ* cells, the loss of *ATG2* caused rapamycin resistance, and over-expression of any of the α-arrestins tested, except those lacking their ^L^/_P_PXY motifs, partially resensitized cells to rapamycin (Fig 2B, column 9, Fig 2C, and S2B-C Figs). Atg2 forms a complex with Atg18, which binds PI3P and PI(3,5)P_2_ and regulates vacuole morphology along with autophagy progression [46,80,83]. We found that the loss of *ATG18*, like *ATG2*, conferred rapamycin resistance (Fig 2B, column 10, Fig 2C and S2B-C Figs). Over-expression of any α-arrestin modestly reduced the rapamycin resistance of *atg18*Δ cells, with Aly1 producing the most robust effect (Figs 2B-C and S2B-C Figs). The Aly K-to-R mutants that block mono-ubiquitination of *ALY*s had more modest effects than their cognate WT alleles.

We examined proteins involved in <u>c</u>ytoplasm-to-<u>v</u>acuole targeting (Cvt), a constitutive form of cargo selective autophagy that utilizes specialized cargo receptors (Atg11, Atg19, and Atg34) [84–93]. APs then deliver Cvt- and receptor-linked cargo to the vacuole [86,91–93]. The loss of *ATG11* or *ATG34* cargo receptors prevented Aly-mediated resistance to rapamycin, in much the same way as occurred in *atg17Δ* cells, while *atg19Δ* had little impact on α-arrestin-induced rapamycin resistance (Fig 2B, columns 15-17, and Fig 2C).

While there were robust genetic interactions with α-arrestins and the autophagy machinery, rapamycin resistance was unaltered when the following genes were deleted: i) *ATG9*, involved in AP isolation membrane seeding and anchoring the Atg2/18 complex to the phagophore membrane [49,56,57], ii) *ATG41*, an Atg9 effector [94], iii) Atg8, a ubiquitin-like protein in autophagy [66,95–97], iv) *ATG5* and *ATG12*, the machinery involved in Atg8 conjugation to autophagic membranes [97–100], or v) *ATG22*, an amino acid permease responsible for the efflux of free amino acids from the vacuole (Figs 2B-C) [101]. Thus, there is a complex network of genetic interactions between the α-arrestins and the autophagic machinery. The genetic interactions between α-arrestins and autophagy initiation, PI3K complex formation and Atg2/Atg18 lipid transfer proteins were the most robust, hinting that α-arrestins may be more closely linked to these stages of autophagy.

### α-Arrestins are required for efficient AP formation and vacuole delivery

In addition to conferring resistance when over-expressed, loss of select α-arrestins increased rapamycin sensitivity (S2D Fig). While the loss of *ART1* was the only single α-arrestin deletion to confer strong sensitivity to rapamycin, the double deletion of the paralogous α-arrestins *BUL1* and *BUL2* also conferred strong sensitivity (S2D Fig). Loss of both *ALY1* and *ALY2* or *RIM8* resulted in a slight increase in sensitivity, consistent with earlier reports for *aly1*Δ *aly2*Δ cells (S2D Fig) [21]. Cells lacking nine of the 14 α-arrestins (9ArrΔ, see S1 Table for genotype) were exquisitely sensitive to rapamycin; this sensitivity must be reliant upon Art1 and Aly1/Aly2 as they are the only α-arrestins missing in this strain background that confer sensitivity to rapamycin on their own.

Based on these findings and the many genetic connections to autophagy revealed by our screen, we decided to assess AP formation and delivery to the vacuole using electron microscopy. To visualize autophagic body (AB) accumulation in the vacuole, we deleted the master vacuolar protease *PEP4*, whose loss severely impairs vacuolar degradation by preventing maturation of vacuolar proteases [102, 103]. Autophagic bodies, which are the remnants of APs after their delivery into the vacuole lumen, are inefficiently degraded in *pep4*Δ cells, thus allowing us to determine how many APs get delivered to the vacuole [104]. Our most severe 9ArrΔ mutant cells accumulated no vacuolar ABs in most cells, but when they were present the ABs were both smaller and less abundant in cells lacking α-arrestins than they were in WT cells (Figs 3A-C and S3A Fig). In the more modest *aly1Δ aly2Δ* or *art1*Δ cells the number of autophagic bodies was not significantly reduced but there was a significant reduction in AB size (Figs 3A-C). As expected, cells lacking *ATG8*, an essential AP component, failed to accumulate any ABs [95].

**Fig 3.**
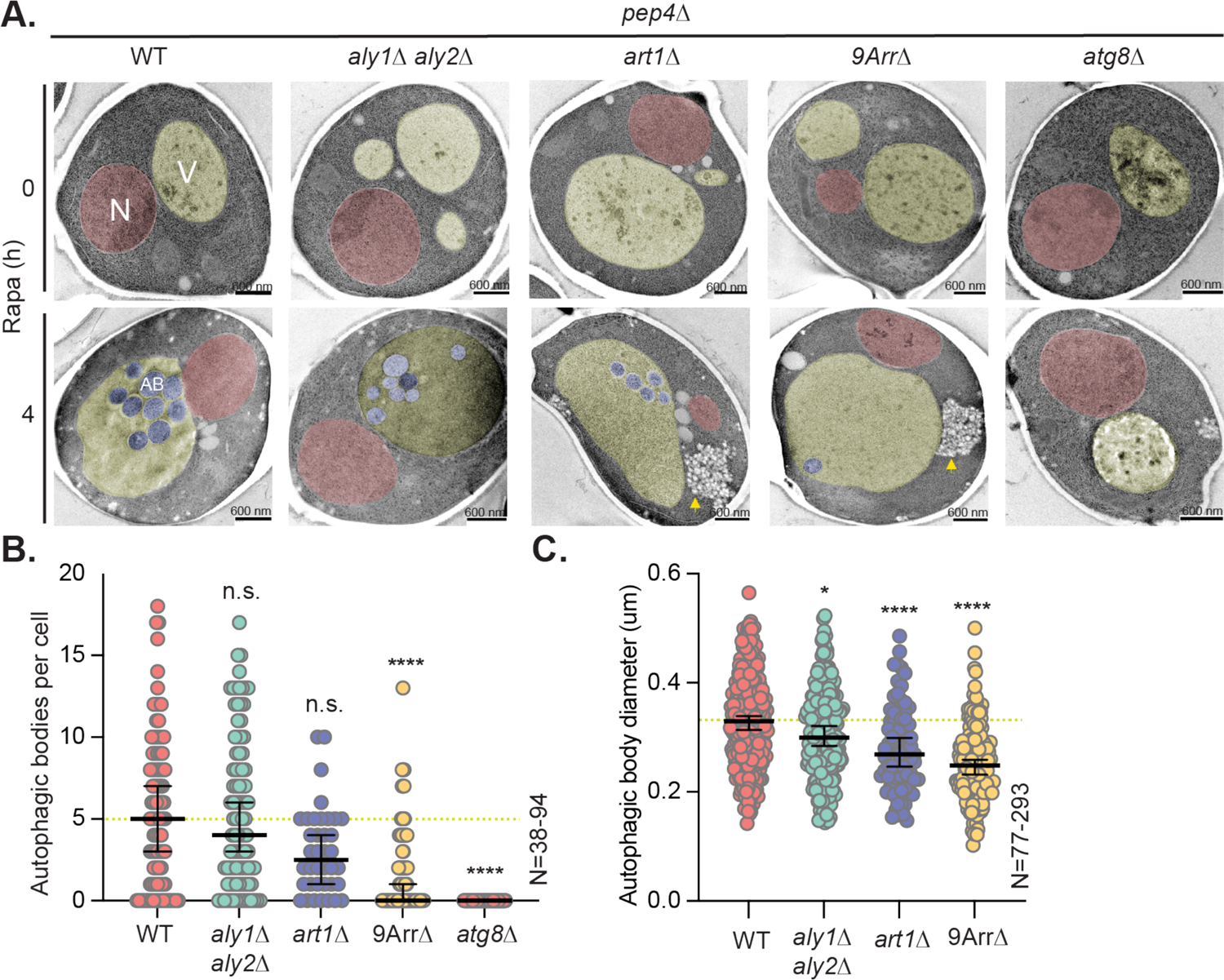
α-Arrestins help promote AP formation and are needed for accumulation of vacuolar autophagic bodies. (**A**) Electron micrographs of cells lacking *PEP4* and the indicated α-arrestins were obtained. Cells are either untreated or treated for 4 h with 200ng/mL rapamycin. Cellular compartments are false colored (N = nucleus in red; V = vacuole in yellow; AB = autophagic body in blue). Yellow arrows mark aberrant structures with low electron density. Scale bars are equal to 600nm. (B-C) The number of autophagic bodies (B) accumulating in the vacuole and diameter of each autophagic body (C) was determined from the EM images in (A) and is plotted. The median number is shown as a black line for each group and error bars represent the 95% confidence interval. For reference, a yellow dashed line represents the median ratio value for WT prior to rapamycin treatment. Kruskal-Wallis statistical analysis with Dunn’s post hoc test was performed to compare each distribution to that of the *pep4*Δ cells) (n.s. = not significant; **** = p-value <0.0001).

We reasoned that perhaps the reduced AB number in α-arrestin mutant cells reflects a defect in AP-vacuole fusion. However, in contrast to cells lacking *VAM3,* a syntaxin-like t-SNARE required for AP-vacuole fusion, there was no dramatic accumulation of cytosolic APs in α-arrestin mutant cells (Fig 3A compared to S3B-C Fig) [105]. These results support the idea that α-arrestins facilitate efficient production and delivery of APs to the vacuole. In addition to alterations in APs, we noticed an increase in low-electron density structures that appeared to emanate from the cortical ER in rapamycin-treated *art1Δ* and *9arrΔ* cells (Fig 3A, yellow arrows). Given their similar electron density to lipids, we posit that these structures represent aberrant lipid accumulations, though their exact nature remains undefined.

To further assess the potential defect in AP delivery to the vacuole, we utilized time-lapse confocal microscopy to capture images of cells expressing plasmid-borne *ATG8pr*-GFP-*ATG8* at short time intervals (every 30s). We then measured the length of time GFP-Atg8 puncta persisted before merging with the vacuole (Fig 4A-B, S1-4 Movies). Atg8, an autophagy-specific ubiquitin-like protein, is conjugated to PE and incorporated into both the inner and outer membranes of APs by Atg2-Atg5 (Fig 2C) [43,66,95,97]. In agreement with our EM results, we found that GFP-Atg8 puncta persisted for longer periods of time in *aly1Δ aly2Δ*, *art1Δ* and 9arrΔ cells than in WT cells (Fig 4A-B). These results indicate a delay in isolation membrane expansion and/or completion, stalling formation of mature APs.

**Fig 4.**
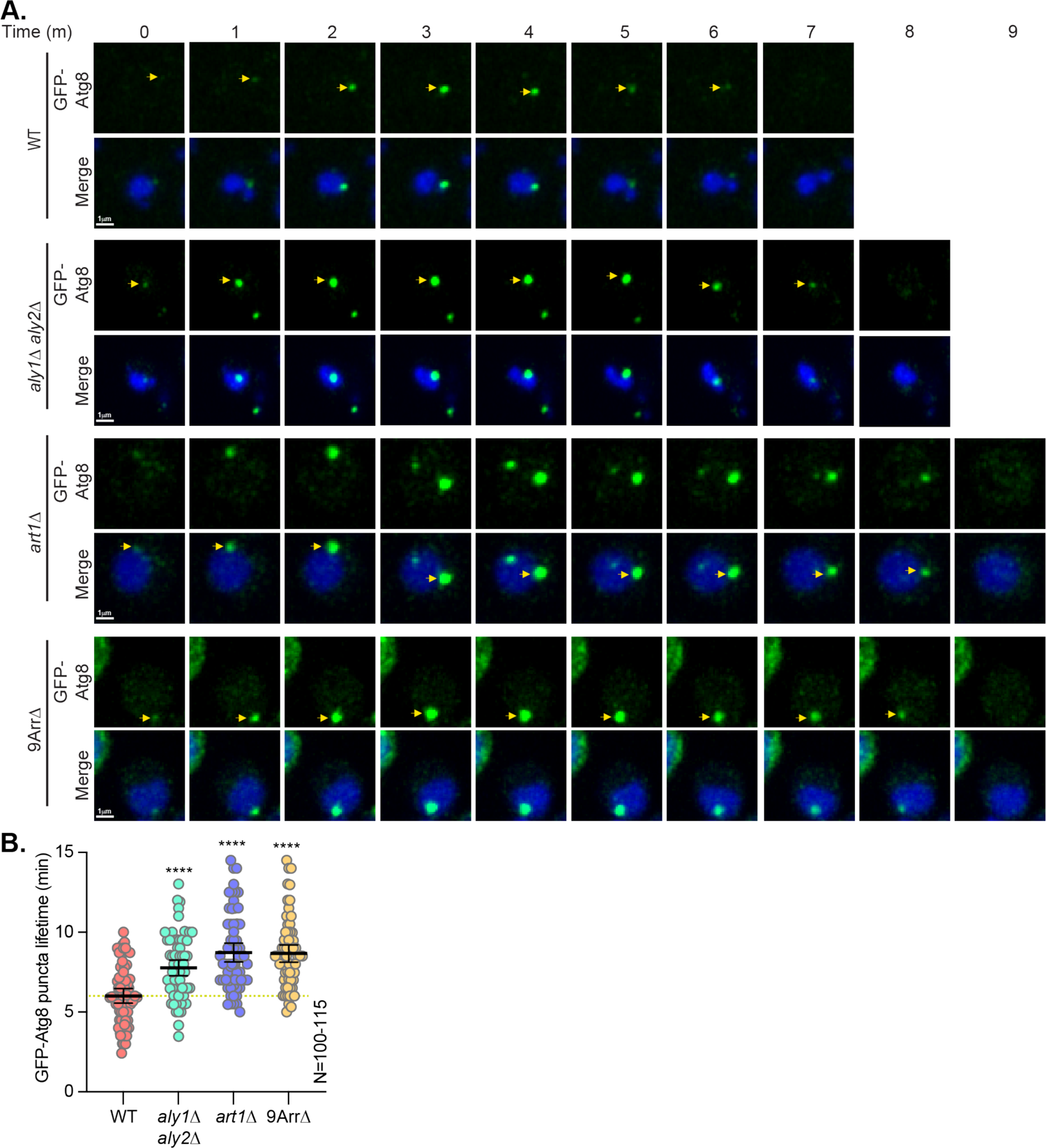
α-Arrestins are required for efficient AP delivery to the vacuole. (**A**) Cells expressing *ATG8pr-*GFP-Atg8 in either WT or α-arrestin mutant cells were imaged by confocal microscopy at 1min intervals following 1h treatment with 200ng/mL rapamycin. CMAC is used to stain the vacuoles (shown in blue). Yellow arrows indicate the GFP-Atg8 puncta, representing APs. 3D projection videos of these cells over the same time periods are available in S1-4 Movies. (B) Quantification of lifetimes for the GFP-Atg8 puncta shown in (A), measured from appearance to merging with the vacuole, is shown as a graph. The lifetime of each puncta is plotted as a circle. The median lifetime is shown as a black line and error bars represent the 95% confidence interval. For reference, a yellow dashed line represents the median lifetime for WT. Kruskal-Wallis statistical analysis with Dunn’s post hoc test was performed to compare the lifetime distributions. Statistical comparisons between mutant and WT cells at that same time point are displayed in black asterisks (n.s. = not significant; four symbols has a p-value <0.0001).

### α-Arrestins are required for efficient GFP-Atg8 autophagic flux

We sought to determine if α-arrestins influence AP formation and delivery by monitoring GFP-Atg8 in cells (Fig 5) [66,95,96]. Prior to TORC1 inhibition, GFP-Atg8 was cytosolically diffuse with rare puncta that likely corresponded to APs from the Cvt pathway (Fig 5A). When TORC1 was inhibited, GFP-Atg8 formed large puncta consistent with PAS recruitment (Fig 5A) [44,106–108]. Once complete, APs fuse with the vacuole and the Atg8 on the outer autophagosomal membrane is cleaved from its PE anchor, releasing it into the cytosol. The Atg8 incorporated into the inner AP membrane is delivered to the vacuole lumen and degraded (Fig 2C) [96]. Inside the vacuole, GFP is cleaved from GFP-Atg8 by vacuolar proteases but is itself degradation resistant. This results in a build-up of free GFP in the vacuole that increases proportionally with the number of APs delivered [109]. Prior to rapamycin treatment, GFP-Atg8 vacuole:whole cell fluorescence ratio was not significantly different in WT and α-arrestin mutants (Fig 5A-B). Upon rapamycin-induced autophagy, α-arrestin mutants had a reduced GFP-Atg8 vacuole:whole cell florescence ratio compared to WT cells. In fact, at 1-hour post rapamycin, all the α-arrestins mutants had failed to increase the vacuole-to-whole cell fluorescence ratio significantly in comparison to their untreated controls, indicative of a defect in autophagic flux in cells lacking α-arrestins (Fig 5B). In contrast, WT cells had significantly elevated GFP-Atg8 vacuole:whole cell florescence ratios at 1-hour post rapamycin and this further increased at 3-hours post rapamycin (Fig 5A-B). At 3 hours post rapamycin, both *art1Δ* and 9ArrΔ cells had significantly reduced vacuole:whole cell fluorescence ratios compared to WT cells. In contrast, *aly1*Δ *aly2*Δ vacuole:whole cell fluorescence ratios were not statistically different from WT yeast at the 3 hour point, suggesting a less severe delay in these mutants. These data support the idea that cells lacking α-arrestins are defective in AP production and/or autophagic flux.

**Fig 5.**
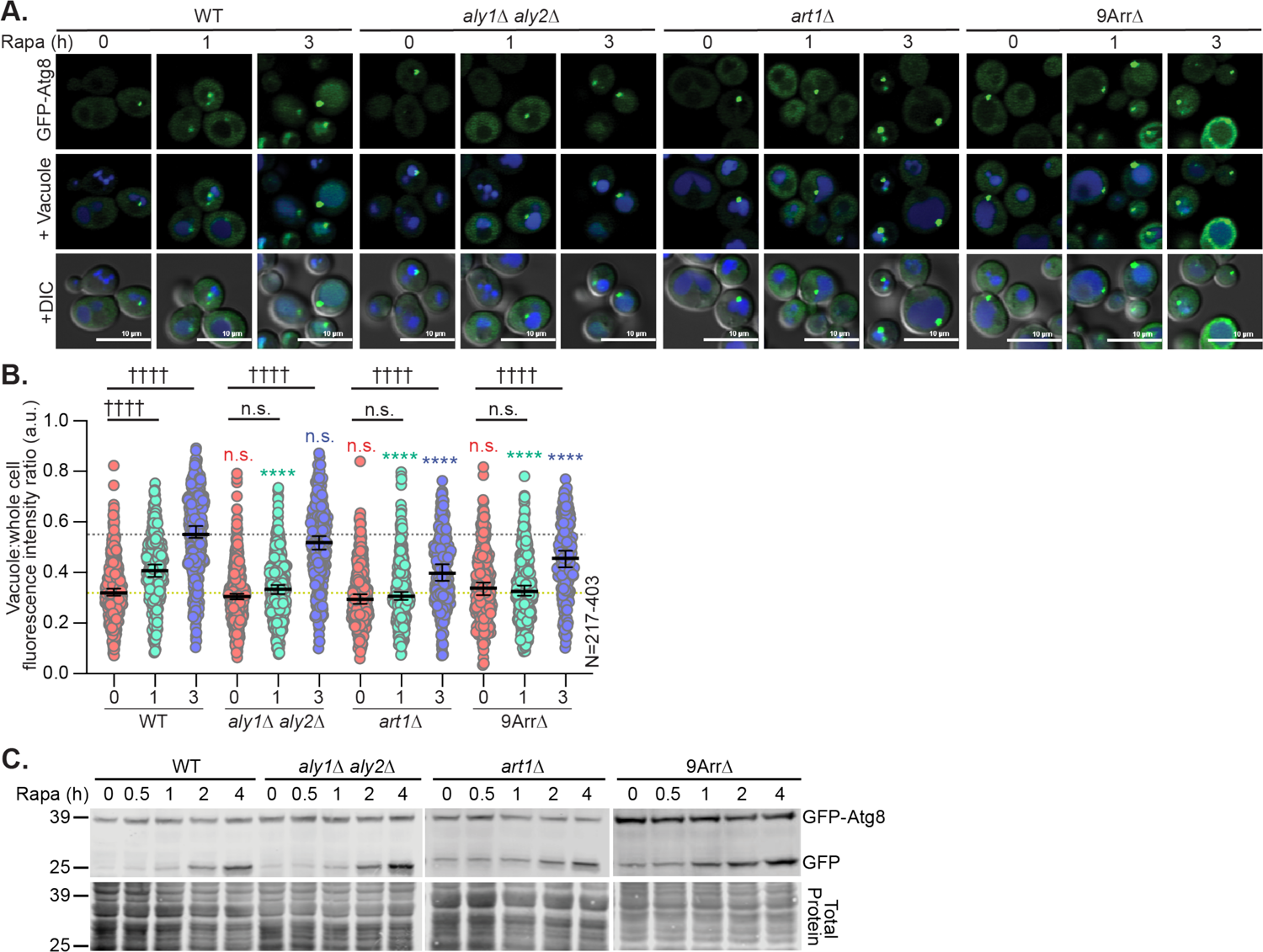
α-Arrestins are required for optimal autophagic flux. (**A**) Cells expressing pRS416-*ATG8pr-*GFP-Atg8 in either WT or α-arrestin mutant cells were imaged by confocal microscopy at the indicated time post-treatment with 200ng/mL rapamycin. CMAC is used to stain the vacuoles (shown in blue). The merge (+DIC) contains the bright field cell image for reference. (B) Quantification of vacuolar to whole-cell fluorescence intensity ratio using Nikon.*ai* software (see *Materials and Methods*) for the cells imaged in panel (A) is shown as a scatter plot. The ratio of vacuolar to whole-cell GFP intensity is plotted for each cell as a circle. The median ratio is shown as a black line and the error bars represent the 95% confidence interval. For reference, yellow and grey dashed lines represent the median ratio value for WT prior to and post rapamycin treatment, respectively. Kruskal-Wallis statistical analysis with Dunn’s post hoc test was performed to compare the ratio distributions. Statistical comparisons between mutant and WT cells at that 0, 1, and 3 hours post rapamycin are shown as red, green, and blue asterisks respectively. Statistical comparisons of WT or mutant cells to their respective t=0 timepoints are shown as black daggers. (n.s. = not significant; four symbols has a p-value <0.0001). (C) Protein extracts from cells as described in (A) were immunoblotted and probed with anti-GFP antibody. The REVERT total protein stain of the membrane is the loading control. Molecular weights are shown on the left side in kDa.

We also assessed GFP-Atg8 degradation in the vacuole using SDS-PAGE and immunoblotting by comparing the abundance of full-length GFP-Atg8 to the free GFP produced after vacuolar cleavage (Fig 5C). Cells lacking α-arrestins had higher GFP-Atg8 levels prior to rapamycin treatment, especially in the 9arrΔ mutant. The free GFP abundance was also increased prior to autophagy induction, suggesting that cells lacking α-arrestins may have higher Atg8 expression, perhaps due to metabolite imbalances in these cells. Following rapamycin treatment, WT cells displayed a more rapid increase in free GFP relative to the intact GFP-Atg8 levels (Fig 5C). In the 9ArrΔ cells, there was increased GFP-Atg8 even before rapamycin treatment. When this increase in GFP-Atg8 abundance was accounted for, the amount of the free GFP was diminished in 9ArrΔ cells relative to the intact GFP-Atg8 in comparison to the WT control (Fig 5C). A more modest, but similar trend, was observed in the *art1*Δ cells but no obvious change was seen in *aly1*Δ *aly2*Δ cells using this approach.

### α-Arrestins permit efficient phagophore expansion

To begin producing isolation membranes for AP production, Atg9, the sole transmembrane protein of the autophagic machinery, is recruited to the PAS. Atg9 brings the membrane in which it resides to serve as ‘seed’ membrane for APs and provides lipid scramblase activity in the growing phagophore (Fig 2C) [56,110–113]. Defective Atg9 recruitment results in impaired AP production [57]. To assess Atg9 abundance and localization, we examined the localization of endogenously tagged Atg9-mNG via live-cell confocal microscopy in WT and α-arrestin mutant cells before and after treatment with rapamycin (S4 Fig). We observed a similar whole-cell fluorescence abundance for Atg9-mNG in WT and α-arrestin mutant cells, and Atg9-mNG abundance increased in all cell types tested post rapamycin treatment (S4B-C Fig). While Atg9 abundance appeared unaffected, we asked if trafficking was disrupted as aberrant Atg9 recycling is associated with defective AP initiation. Atg9 recruitment to the PAS and recycling is dependent on the clathrin adaptor complex, AP-3, and retromer, which aids in localizing Atg9’s cofactor Atg27 [113, 114]. We found that two AP-3-dependent cargo, Ypq2 and Yck3, had normal vacuolar membrane localization in α-arrestin mutants, suggesting that AP-3 trafficking was not disrupted in α-arrestin mutant cells (S5A-B Fig) [115, 116]. Similarly, neither retromer subunit, Vps17, nor its cargo, Vps10, were mislocalized in α-arrestin mutants, suggesting that the trafficking pathways controlling Atg9 remained robust when α-arrestins were absent (S5C-D Fig) [59, 117].

To further probe Atg9 dynamics, we made use of *atg8*Δ cells. In the absence of *ATG8*, AP production is stalled at the PAS, but Atg9 and Ape1, an autophagic cargo in the Cvt pathway, continue to be recruited to try to stimulate AP formation [118]. We used mCherry-tagged Ape1 to mark the PAS in *atg8Δ* cells where AP formation was stalled. In cells lacking *ATG1*, a negative control in these experiments that prevents initiation of autophagy, Atg9 fails to co-localize with the PAS after rapamycin treatment (Fig 6A) [57]. In contrast, loss of α-arrestins did not significantly change the amount of Atg9 co-localized to the PAS in comparison to WT cells (Fig 6A-B). Taken together, our evaluations of Atg9 and its related trafficking pathway suggest that the α-arrestin-associated impairment of autophagy was not the result of defective Atg9 trafficking or PAS recruitment.

**Fig 6.**
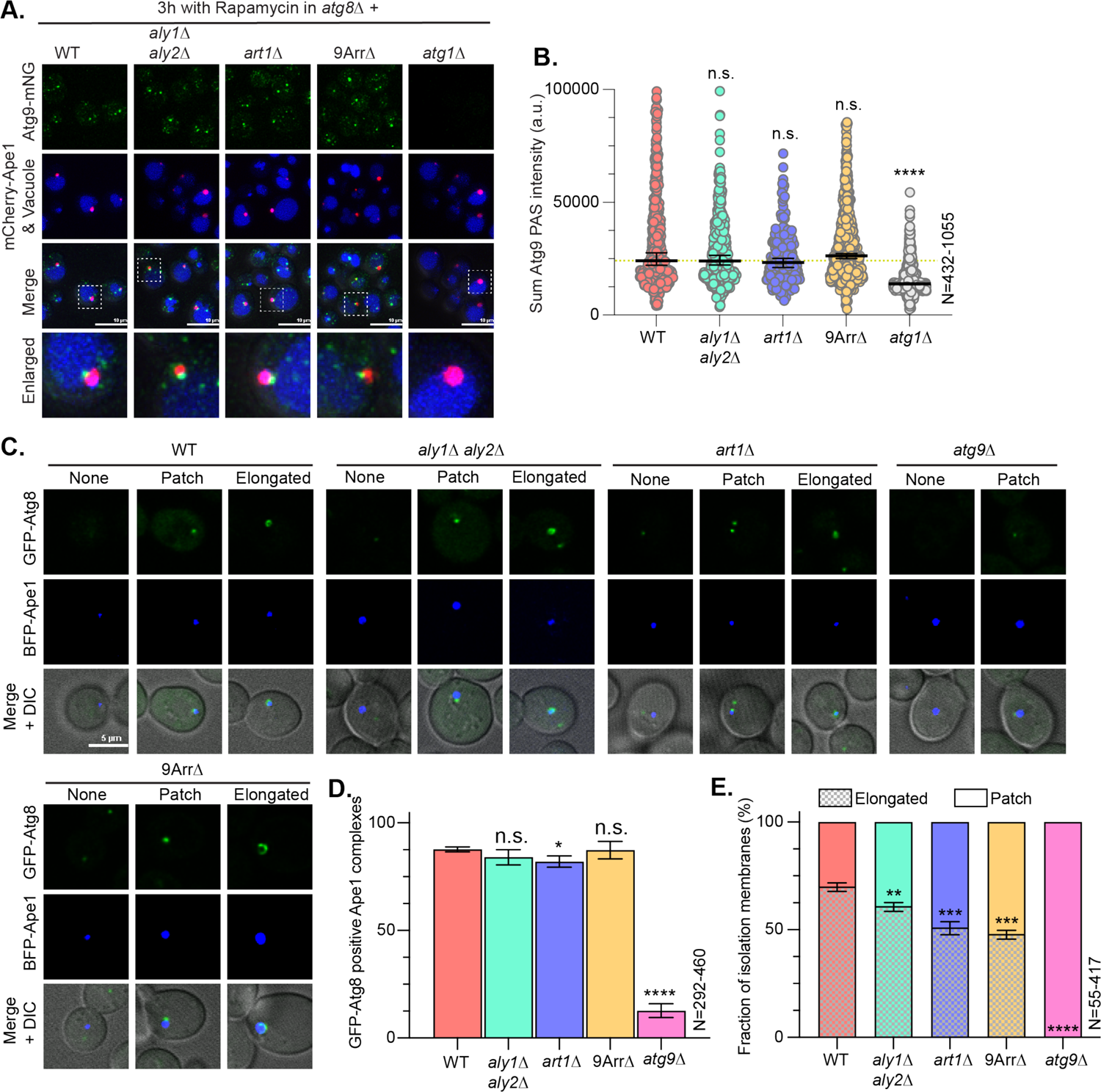
α-Arrestins are required for proper phagophore expansion. (**A**) Cells lacking *ATG8*, which stalls AP production, and expressing chromosomally integrated Atg9-mNG from its endogenous locus in either WT, *atg1Δ,* or α-arrestin mutant, were imaged by confocal microscopy after 3h treatment with 200ng/mL rapamycin. pRS416-*ADH1pr-* mCherry-Ape1 (gift from Segev lab, Univ. of Illinois at Chicago) was used to mark the PAS, while CMAC and FM4-64 were used to stain the vacuolar lumen (shown in blue) or membrane (shown in red), respectively. (B) The sum of the Atg9-mNG intensity at mCherry-Ape1-marked puncta, which represents the PAS, using Nikon.*ai* software (see methods) for the cells imaged as in (A) was plotted for each cell as a circle. The median value is shown as a black line and the error bars represent the 95% confidence interval. For reference, a yellow dashed line represents the median intensity value for WT. Kruskal-Wallis statistical analysis with Dunn’s post hoc test was performed to compare the intensity distributions. Statistical comparisons between mutant and WT cells are displayed in black asterisks (n.s. = not significant; **** = p-value <0.0001). (C) Representative images displaying Atg8-positive structures from cells expressing *ATG8pr*-GFP-Atg8 [2] under the control of its own promoter and *CUP1pr*-BFP-Ape1 (gift from Kraft Lab, Univ. of Freiburg) were imaged by confocal microscopy 1h post-treatment with 200ng/mL rapamycin. The merge (+DIC) contains the bright field cell image for reference. (D) The mean percentage of cells with GFP-Atg8 positive Ape1 puncta, indicative of nucleated APs, from the images in (C) were plotted. Error bars represent the standard deviation. Unpaired t tests were performed to compare the percentages of GFP-Atg8 positive Ape1 puncta. Statistical comparisons between mutant and WT cells are displayed in black asterisks (n.s. = not significant; **** = p-value <0.0001). (E) The mean percentage of cells with either elongated GFP-Atg8 (colored bar), indicative of AP membrane expansion, or patched GFP-Atg8 (colored and hatched bar), indicative of initiated but not expanded APs, from the images in (C) were plotted. Error bars represent the standard deviation. Unpaired t tests were performed to compare the percentages of elongated structures. Statistical comparisons between mutant and WT cells are displayed in black asterisks (n.s. = not significant; **** = p-value <0.0001).

Next, to monitor phagophore expansion, we turned to the ‘giant Ape1’ phagophore expansion assay [57]. In this assay, Ape1 is over-expressed and forms a massive complex (aka ‘giant’ Ape1) at the PAS as the autophagic machinery attempts to expand the isolation membrane around this cargo [57]. By over-expressing BFP-tagged Ape1 with GFP-Atg8, we assessed the ability of WT and α-arrestin mutant cells to recruit Atg8 and expand the isolation membrane. During this process, GFP-Atg8 forms a tell-tale, cup-like shape around the BFP-tagged Ape1 as it engulfs the expansive cargo (Fig 6C-E). We observed no change in the ability of cells lacking α-arrestins to recruit GFP-Atg8 to giant Ape1 structures in comparison to WT cells (Fig 6D), further demonstrating that Atg9 recruitment and isolation membrane seeding were not impaired in α-arrestin mutants (Figs 6A-B, S4 Fig).

In contrast to these data, we noted that membrane seeding was severely impaired and little to no GFP-Atg8 was recruited to the Ape1 patches in our *atg9*Δ control cells (Fig 6C-E) [57]. Though Atg9 recruitment was normal, in cells lacking α-arrestins fewer of the cup-like GFP-Atg8 structures associated with autophagophore membrane expansion formed (Fig 6E). Thus, phagophore expansion, as measured by GFP-Atg8 encapsulation of Ape1, was significantly reduced in *aly1Δ aly2Δ* cells, and further delayed in *art1Δ* or 9arrΔ cells. We propose that cells lacking α-arrestins are defective in phagophore expansion, consistent with the prolonged GFP-Atg8 lifetimes and reduced AB accumulation we observed in other assays (Figs 3 and 4).

### TORC1 localization and Atg13 regulation are unaffected by α-arrestins

The TORC1 signaling complex inhibits autophagy under nutrient rich conditions by phosphorylating Atg13, which thereby prevents Atg1 kinase activation and downstream events (Fig 2C) [5, 6]. Given the defective AP delivery and genetic interactions between α-arrestins and the autophagy initiation complex, we next assessed the phospho-status of endogenous Atg13 by examining its electrophoretic mobility (Fig 7A). We observed a similar electrophoretic profile for Atg13 in cells lacking α-arrestins or WT cells prior to rapamycin treatment, consistent with TORC1 phosphorylation and inhibition of Atg13. Following addition of rapamycin, Atg13 increased in electrophoretic mobility in both WT cells and those lacking α-arrestins, as would be expected when TORC1-mediated phosphorylation is inhibited and Atg13 is activated by dephosphorylation (Fig 7A) [6]. It appears that cells lacking α-arrestins can activate Atg13 as well as WT cells in response to rapamycin.

**Fig 7.**
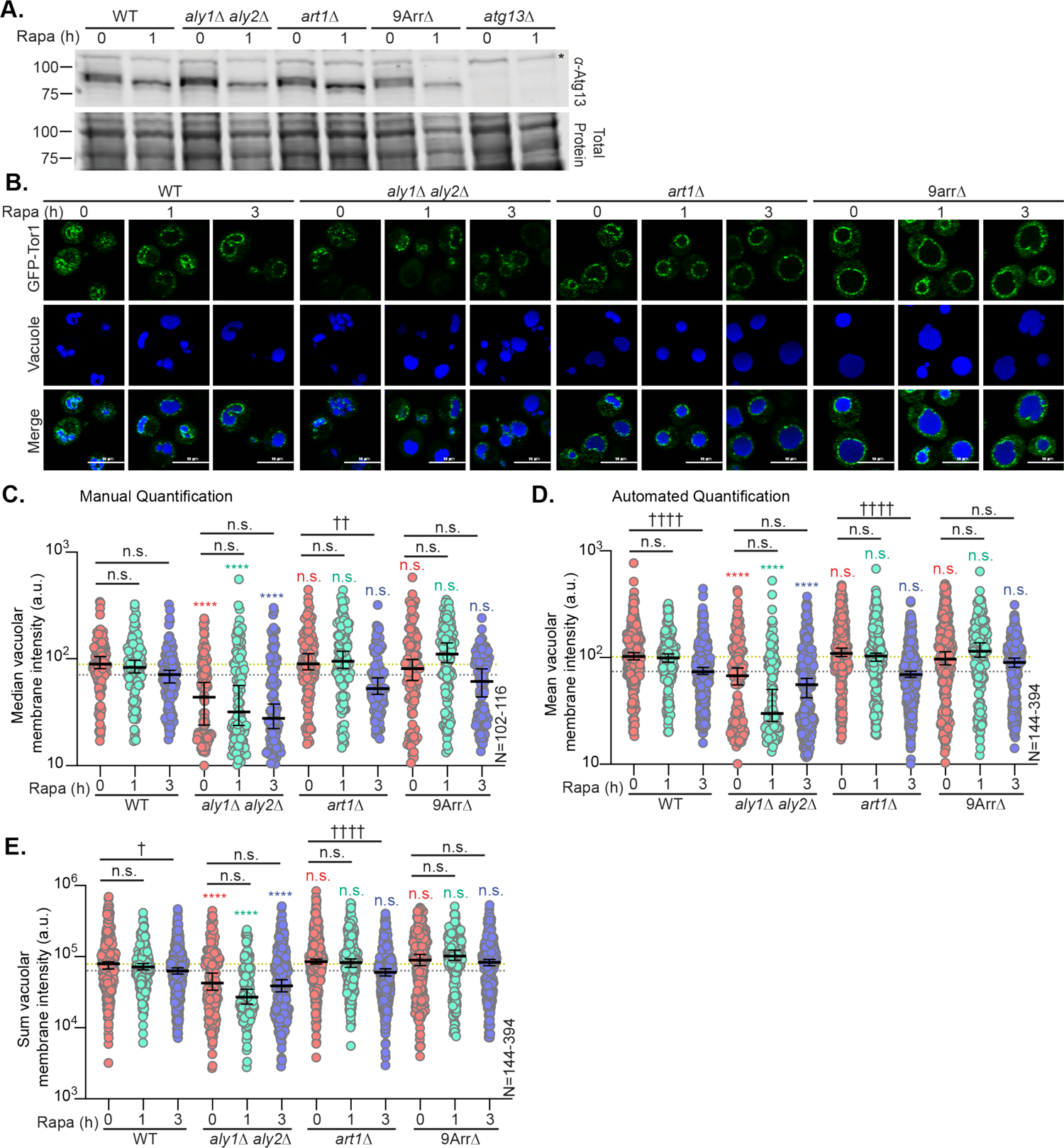
Tor1 signaling and localization are not affected by the loss of α-arrestins. (**A**) WT cells or cells lacking the indicated α-arrestin were harvested at the indicated time points post-treatment with 200ng/mL rapamycin. Whole-cell protein extracts were made, analyzed by SDS-PAGE, immunoblotted and probed with anti-Atg13 antibody (gift from Klionsky lab, Univ. of Michigan). REVERT total protein stain of the membrane is the loading control and molecular weights are shown on the left in kDa. A non-specific band is marked with an asterisk (*). (B) Cells expressing *TOR1pr*-Tor1-GFP (gift from Ford lab, Univ. of Pittsburgh) in either WT or α-arrestin mutant cells were imaged by confocal microscopy at the indicated times post-treatment with 200ng/mL rapamycin. CMAC is used to stain the vacuoles (shown in blue). (C) Manual quantification of the median GFP intensity at the vacuolar membrane using ImageJ software (see *Materials and Methods*) for the cells in panel (B) is shown as a graph. The median GFP intensity is plotted for each cell as a circle. The median value of each population is shown as a black line and the error bars represent the 95% confidence interval. For reference, a yellow or grey dashed line represents the median value for WT prior to or after 3h of rapamycin treatment, respectively. Kruskal-Wallis statistical analysis with Dunn’s post hoc test was performed to compare the intensity distributions. Statistical comparisons between mutant and WT cells at that 0, 1, and 3 hours post rapamycin are shown as red, green, and blue asterisks, respectively. Statistical comparisons of WT or mutant cells to their respective t=0 timepoints are shown as black daggers/text. (n.s. = not significant; four symbols has a p-value <0.0001). (D) Quantification of the mean GFP intensity at the vacuolar membrane using Nikon.*ai* software (see methods) for the cells in panel (B) is shown as a graph. The mean GFP intensity is plotted for each cell as a circle. The median value is shown as a black line and the error bars represent the 95% confidence interval. For reference, a yellow or grey dashed line represents the median value for WT prior to or after 3h of rapamycin treatment, respectively. Statistical analyses and comparisons are as in (C). (E) Quantification of the sum GFP intensity at the vacuolar membrane using Nikon.*ai* software for the cells in panel (A) is shown as a graph. The sum GFP intensity is plotted for each cell as a circle. The median value is shown as a black line and the error bars represent the 95% confidence interval. For reference, a yellow or grey dashed line represents the median value for WT prior to or after 3h of rapamycin treatment, respectively. Statistical analyses and comparisons are as in (C).

While predominantly localized to the vacuolar membrane, there is a second population of TORC1 residing on endosomes that is responsible for control of autophagy through phosphorylation of Atg13 [3, 119]. We examined the localization and abundance of GFP-Tor1, the kinase subunit in the TORC1 complex, using live-cell confocal microscopy and quantified GFP-Tor1 intensities at the vacuole membrane using two approaches (Fig 7B-E). In WT cells and those lacking α-arrestins, Tor1 appeared robustly localized to the vacuole membrane both before and after rapamycin treatment (Fig 7B). We evaluated Tor1 vacuolar membrane intensity using a manual quantification protocol (see *Materials and Methods*) and an automated quantification pipeline using the machine learning platform NIS.*ai*. We found strong agreement between the two methods, supporting the use of our automated method (compare Figs 7C-D). There was no significant difference in the vacuolar localization of GFP-Tor1 between WT cells and those lacking α-arrestin prior to rapamycin addition based on the median vacuolar fluorescence intensity measurements (Figs 7B-D). However, we noted that the vacuole morphology and size was dramatically different between WT cells and those lacking α-arrestins; cells lacking Alys had hyper-fragmented vacuoles and those lacking Art1 or the 9ArrΔ cells had enlarged, unilobed vacuoles (Fig 7B and S6A Fig). The length of the vacuole membrane itself was significantly larger in *art1*Δ and 9ArrΔ cells than in WT cells prior to rapamycin treatment (S6B-C Fig).

As a result of vacuole membrane length changes, we felt it was important to not only assess the concentration of fluorescently tagged proteins, as indicated by the mean GFP pixel intensity at this organelle, but also their overall abundance, as determined by summing the pixel intensities along the vacuole membrane. Given the vacuole morphology changes in α-arrestin mutants, we apply this kind of analysis throughout when assessing vacuole membrane localized proteins. Using this approach, we found that in WT cells the concentration of GFP-Tor1 decreased at the vacuolar membrane following rapamycin treatment (Fig 7C-D), but the length of the vacuole membrane also increased (S6B-C Fig) in response to this treatment. Therefore, the overall amount of Tor1 at the vacuole was only slightly diminished in response to rapamycin (Fig 7E). Thus, the observed reduction in the mean fluorescence intensity of Tor1 at the vacuole post-rapamycin treatment is more likely because GFP-Tor1 is now distributed across a larger vacuole membrane rather than being the result of reduced Tor1 localization to this organelle (Figs 7C-E and S6B-C Fig). For *art1*Δ cells, Tor1 concentration drops at the vacuole membrane post rapamycin, but this also likely reflects the increase in vacuole membrane area (Figs 7C-D and S6B-C). However, in 9ArrΔ cells, there were no significant changes in the concentration or total amount of Tor1 at the vacuole membrane in response to rapamycin, nor were the changes in vacuole size post-rapamycin as substantial in these cells (Figs 7C-E and S6B-C). Cells lacking Alys appeared to be comprised of two populations in the Tor1 imaging, with one population having dramatically reduced Tor1 concentration and abundance both before and after rapamycin treatment, even though the vacuole area increased in response to rapamycin in the same way as WT cells (Figs 7C-E and S6B-C). This altered Tor1 abundance could contribute to the defective autophagy observed in cells lacking Alys, although if it is linked, it is not through Atg13 regulation (Fig 7A). Since over-expression of α-arrestins causes resistance to rapamycin, we assessed Tor1 localization in WT cells or those over-expressing Aly1, Aly2, or Art1 (S7A-C Fig). We observed no dramatic changes in the density or abundance of Tor1 at the vacuolar membrane in cells over-expressing these α-arrestins, suggesting that this is not a driver for altered rapamycin resistance. These data suggest that Aly2 and Art1 are not dramatically affecting the distribution of Tor1 or TORC1 control of autophagy through Atg13. Aly1 had a more significant influence on Tor1 abundance, but despite this did not seem to alter Atg13 signaling.

### α-Arrestins prevent aberrant Atg2-Atg18 accumulation on the vacuole membrane

Once seeded, isolation membranes expand rapidly with the aid of the Atg18-Atg2 lipid transfer complex, which tethers the PAS to the ER and supplies the growing phagophore with lipids (Fig 2C) [47,50,51]. Since α-arrestin mutants were defective in isolation membrane expansion and had strong genetic interactions with Atg2-Atg18 (Fig 2, S1-2 Figs), we next examined Atg18 and Atg2 localization. Using fluorescently tagged GFP-Atg18, we assessed the stability and the vacuolar membrane abundance of Atg18 (Fig 8). There was no change in Atg18 protein levels, as determined by immunoblotting, between WT cells or those lacking α-arrestins (Fig 8A). Despite similar Atg18 protein levels, by live-cell confocal microscopy we observed a significant increase in both the concentration and abundance of GFP-Atg18 at the vacuolar membrane in *art1Δ* and, more strikingly, 9ArrΔ cells prior to rapamycin addition (Fig 8B-D). While *aly1*Δ *aly2*Δ cells did not have more Atg18 at the vacuole membrane to begin with, after 1 hour of rapamycin treatment Atg18 density on the vacuole membrane (mean fluorescence intensity in Fig 8C) and overall abundance (sum vacuolar intensity in Fig 8D) were significantly higher in cells lacking Alys than they were in WT cells. Post-rapamycin treatment, the Atg18 abundance and concentration remained elevated in 9Arr cells relative to the WT control (Fig 8B-D). For *art1*Δ cells, by 3h post rapamycin the Atg18 abundance and concentration at the vacuole membrane had lessened relative to what it was prior to rapamycin addition (Fig 8B-D), suggesting that the defect in this strain is more modest than that of 9ArrΔ cells. Thus, in cells lacking α-arrestins, Atg18 is aberrantly retained on the limiting membrane of the vacuole.

**Fig 8.**
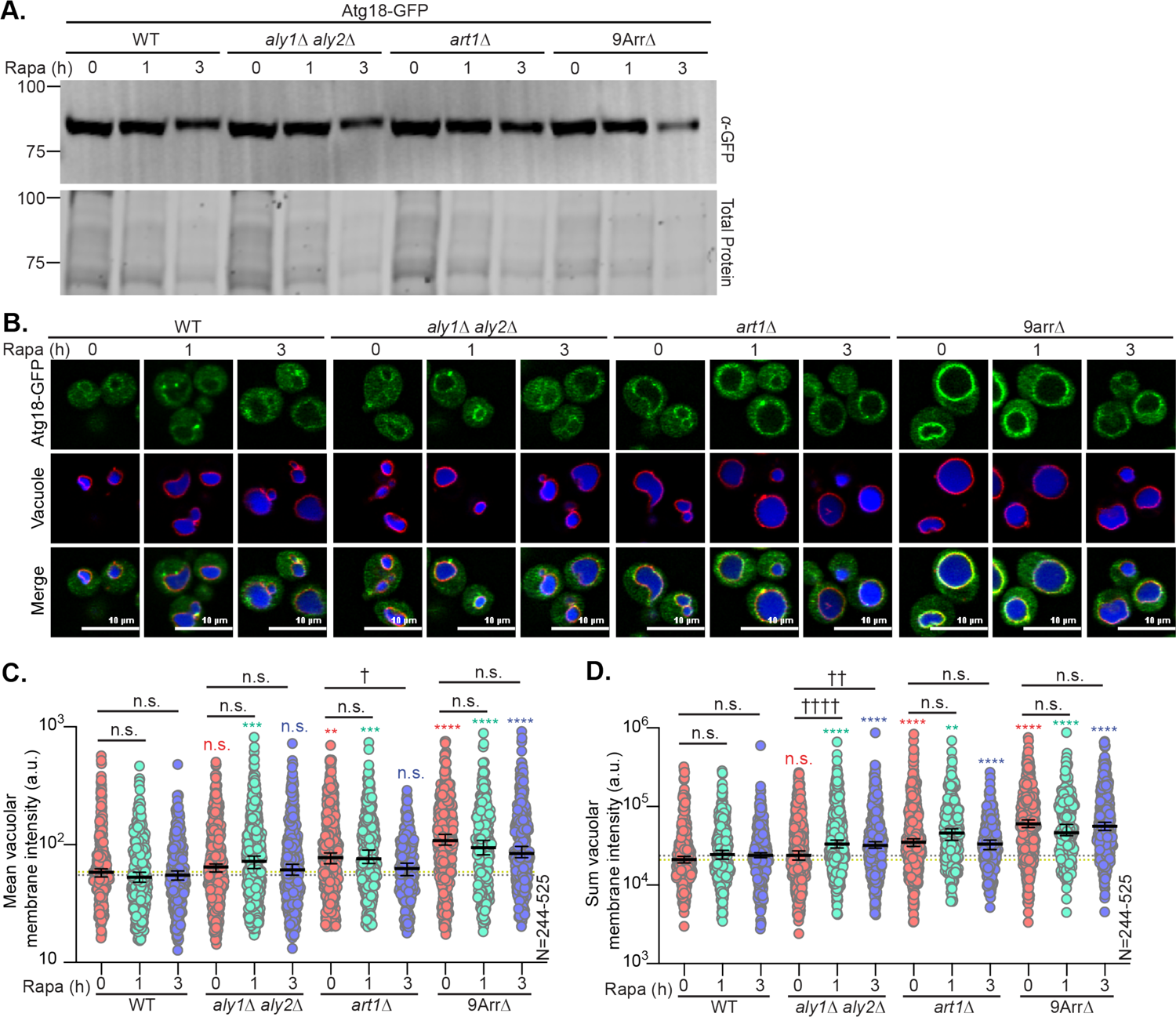
Lipid transfer protein Atg18, required for phagophore expansion, is mislocalized in the absence of α-arrestins. (**A**) Cells expressing *ATG18pr*-Atg18-GFP in either WT or α-arrestin mutant cells were harvested at the indicated time points post-treatment with 200ng/mL rapamycin. Whole-cell protein extracts were made, analyzed by SDS-PAGE, immunoblotted and probed with anti-GFP antibody (Santa Cruz Biotechnology, Santa Cruz, CA). REVERT total protein stain of the membrane is the loading control and molecular weights are shown on the left in kDa. (B) Cells expressing *ATG18pr*-Atg18-GFP (gift from Kraft Lab, Univ. of Freiburg) in either WT or α-arrestin mutant cells were imaged by confocal microscopy at the indicated time post-treatment with 200ng/mL rapamycin. CMAC and FM4-64 is used to stain the vacuolar lumen (shown in blue) and membrane (shown in red), respectively. (C) The mean GFP intensity at the vacuolar membrane using Nikon.*ai* software (see methods) for the cells in (A) is plotted for each cell as a circle. The median value is shown as a black line and the error bars represent the 95% confidence interval. A yellow or grey dashed line represents the median intensity value for WT prior to or following 3h of rapamycin treatment, respectively. Kruskal-Wallis statistical analysis with Dunn’s post hoc test was performed to compare the intensity distributions. Statistical comparisons between mutant and WT cells at that 0, 1, and 3 hours post rapamycin are shown as red, green, and blue asterisks respectively. Statistical comparisons of WT or mutant cells to their respective t=0 timepoints are shown as black daggers. (n.s. = not significant; four symbols has a p-value <0.0001). (D) The sum GFP intensity at the vacuolar membrane using Nikon.*ai* software for the cells in (A) is plotted for each cell as a circle. The median value is shown as a black line and the error bars represent the 95% confidence interval. A yellow or grey dashed line represents the median intensity value for WT prior to or following 3h of rapamycin treatment, respectively. Statistical analyses and comparisons are as in (C).

A similar localization defect was observed for Atg2, a Vps13-family lipid transfer protein that localizes predominantly to ER-associated puncta prior to rapamycin treatment in WT cells and shifts to the vacuolar membrane as they tether to the PAS after autophagy induction (Fig 2C) [49, 50]. The binding of Atg18 to Atg2 is important for the complex’s localization to the PAS [50,120,121]. Like Atg18, we found that α-arrestin mutant cells had increased Atg2 at the vacuolar membrane in basal growth conditions (S8A-C Fig). Though the average intensity of Atg2-GFP at the vacuole membrane in α-arrestin mutants returns to WT levels after treatment with rapamycin (S8B Fig), the total abundance of Atg2 remains higher (S8C Fig). Together, these results indicate that, while α-arrestin mutants can correct for the increased concentration of Atg2 at the vacuolar surface once autophagy is induced, cells lacking α-arrestins are retaining more Atg18 and Atg2 at the vacuole. Given the role for this complex in phagophore expansion, this may provide a functional link to the α-arrestin mutant defects in autophagic body size and number, as well as AP lifetime and expansion (Figs 3-6).

### α-Arrestins help maintain the appropriate balance of phospholipid regulators at the vacuolar membrane

Atg18 is recruited to the PAS through its interactions with PI3P and PI(3,5)P_2_, which are required for Atg18’s role in regulating vacuolar morphology [46,61,83,122–124]. Atg18 regulates the activity of the lone enzyme responsible for the production of PI(3,5)P_2_ in yeast, Fab1 [83,125,126]. Because Atg18 is mislocalized when α-arrestins are absent, we next examined the localization of phospholipid regulators, Vps34 and Fab1, and PI3P in cells lacking α-arrestins. We utilized a Fab1-ENVY construct to determine the localization and vacuolar membrane abundance of the PI(3,5)P_2_ kinase (Fig 9A-C). Like our results with GFP-Atg18, we found increased concentrations (mean fluorescence intensity, Fig 9B) and abundance (sum fluorescence intensity, Fig 9C) of Fab1-ENVY at the vacuolar membrane in *art1Δ* and 9arrΔ cells, both before and after rapamycin addition (Fig 9B-C). However, Fab1-ENVY localization was unchanged in *aly1*Δ *aly2*Δ cells, which have consistently had more modest defects than those observed in the *art1*Δ or 9ArrΔ cells. These results suggest that in the absence of α-arrestins, cells have more Fab1 on the vacuole, which could in turn increase PI(3,5)P_2_ at this location and potentially contribute to the mislocalization of Atg18-Atg2.

**Fig 9.**
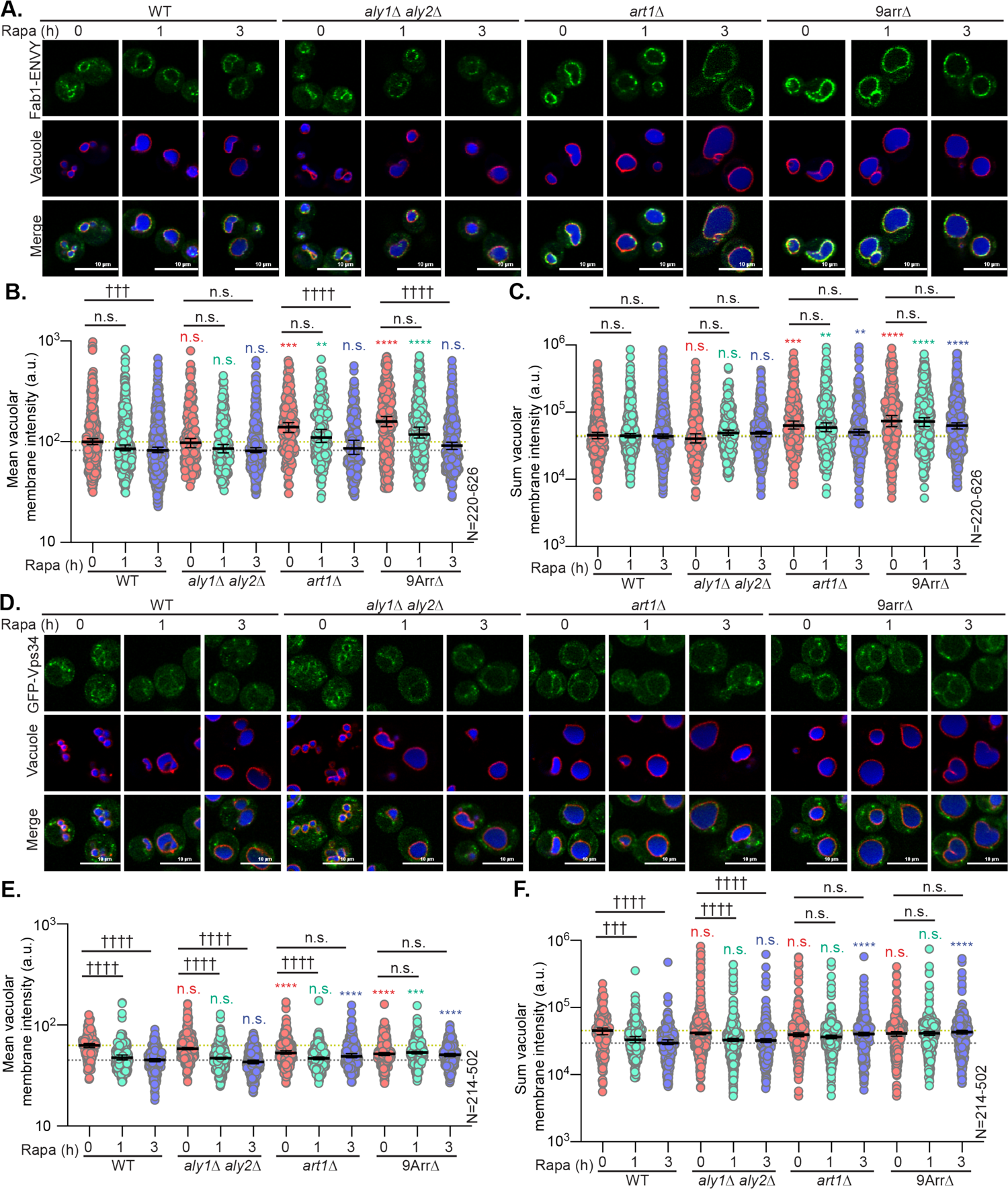
α-Arrestins are required for the proper localization of the phospholipid-modifying enzymes Fab1 and Vps34. (**A**) Cells expressing *FAB1pr*-Fab1-ENVY (gift from Weisman lab, Univ. of Michigan) in either WT or α-arrestin mutant cells were imaged by confocal microscopy at the indicated time post-treatment with 200ng/mL rapamycin. CMAC and FM4-64 was used to stain the vacuolar lumen (shown in blue) or membrane (shown in red), respectively. (B) Quantification of the mean ENVY intensity at the vacuolar membrane using Nikon.*ai* software (see *Materials and Methods*) for the cells in (A) is plotted for each cell as a circle. The median value is shown as a black line and the error bars represent the 95% confidence interval. For reference, a yellow or grey dashed line represents the median intensity value for WT prior to or following 3h of rapamycin treatment, respectively. Kruskal-Wallis statistical analysis with Dunn’s post hoc test was performed to compare the intensity distributions. Statistical comparisons between mutant and WT cells at that 0, 1, and 3 hours post rapamycin are shown as red, green, and blue asterisks respectively. Statistical comparisons of WT or mutant cells to their respective t=0 timepoints are shown as black daggers. (n.s. = not significant; four symbols has a p-value <0.0001). (C) Quantification of the sum of ENVY fluorescence intensity at the vacuolar membrane using Nikon.*ai* software for the cells in (A) is plotted for each cell as a circle. The median value is shown as a black line and the error bars represent the 95% confidence interval. For reference, a yellow or grey dashed line represents the median intensity value for WT prior to or following 3h of rapamycin treatment, respectively. Statistical analyses and comparisons are as in (B). (D) Cells expressing *VPS34pr*-GFP-Vps34 (gift from Weisman lab, Univ. of Michigan) were imaged by confocal microscopy as in (A). (E) Quantification of the mean GFP intensity at the vacuolar membrane using Nikon.*ai* software (see *Materials and Methods*) for the cells in (D) is plotted for each cell as a circle. The description of features is the plot are the same as those indicated in (B). (F) Quantification of the sum GFP intensity at the vacuolar membrane using Nikon.*ai* software for the cells in (D) is plotted for each cell as a circle. Statistical analyses and comparisons are the same as those described in (B).

Atg18, as well as other components of the autophagic machinery are recruited to the PAS by binding PI3P, a precursor to PI(3,5)P_2_ and a key component of the autophagosome isolation membrane [45,126–129]. The autophagic machinery includes a dedicated PI3K complex, called PI3K Complex I [45,77,78]. PI3K Complex I localizes to the vacuolar membrane and the PAS, producing PI3P for the isolation membrane to recruit other components of the autophagic machinery, including the Atg18-Atg2 lipid-transfer complex, to support AP production [47, 127]. While essential for autophagy, hyperactivity of the PI3K complexes’ catalytic subunit, Vps34, inhibits autophagic flux by delaying the release of machinery from completed APs and slowing their fusion to the vacuole [54]. Given the defective localization of Atg18, Atg2, and Fab1 in α-arrestin mutant cells, as well as the genetic interactions between Atg2, Atg18, and the PI3K Complex I and II member, Atg6, with the α-arrestins (see Fig 2), we next examined the localization of the PI3K. We evaluated GFP-Vps34 localization and vacuolar membrane abundance using live-cell confocal microscopy (Fig 9D-F). We found that, while *aly1Δ aly2Δ* had no effect on GFP-Vps34 concentration or abundance at any time point, *art1Δ* and 9AarrΔ had lower levels of Vps34 at the vacuolar membrane under basal conditions (Fig 9D-E). However, unlike WT and *aly1Δ aly2Δ* cells, Vps34 concentration (mean fluorescence intensity, Fig 9E) and overall abundance (sum fluorescence intensity, Fig 9F) was unchanged following rapamycin treatment in *art1Δ* and 9arrΔ (Fig 9D-E). This enrichment of Vps34 in cells lacking α-arrestins could be an indicator of elevated vacuolar PI3P levels.

To assess the sub-cellular distribution of PI3P, we examined the residence of a fluorescently tagged PI3P probe, GFP-FYVE(Eea1) [130]. Typically, PI3P is enriched in endosomal membranes where it’s important for recruiting PI3P binding proteins [58,59,117,131–135]. However, cells lacking α-arrestins had increased GFP-FYVE(Eea1) at the limiting vacuolar membrane compared to WT cells (Fig 10A-B). While puncta adjacent to vacuoles, consistent with endosomal localization, were observed in all cells examined, there was distinct vacuole membrane localization in >40% of cells lacking α-arrestins whereas less than 20% of WT cells had GFP-FYVe on the vacuole membrane (Fig 10B). We suggest a model in which the PI3P imbalance results in mislocalized and retained Atg18-Atg2 on the vacuole membrane in α-arrestin mutants (Fig 10C). We posit that this in turn leads to defects in AP membrane expansion, as Atg18-Atg2 cannot operate at the PAS as they should, increasing the AP lifetime prior to vacuole fusion to delay autophagic flux to the vacuole (Fig 10C).

**Fig 10.**
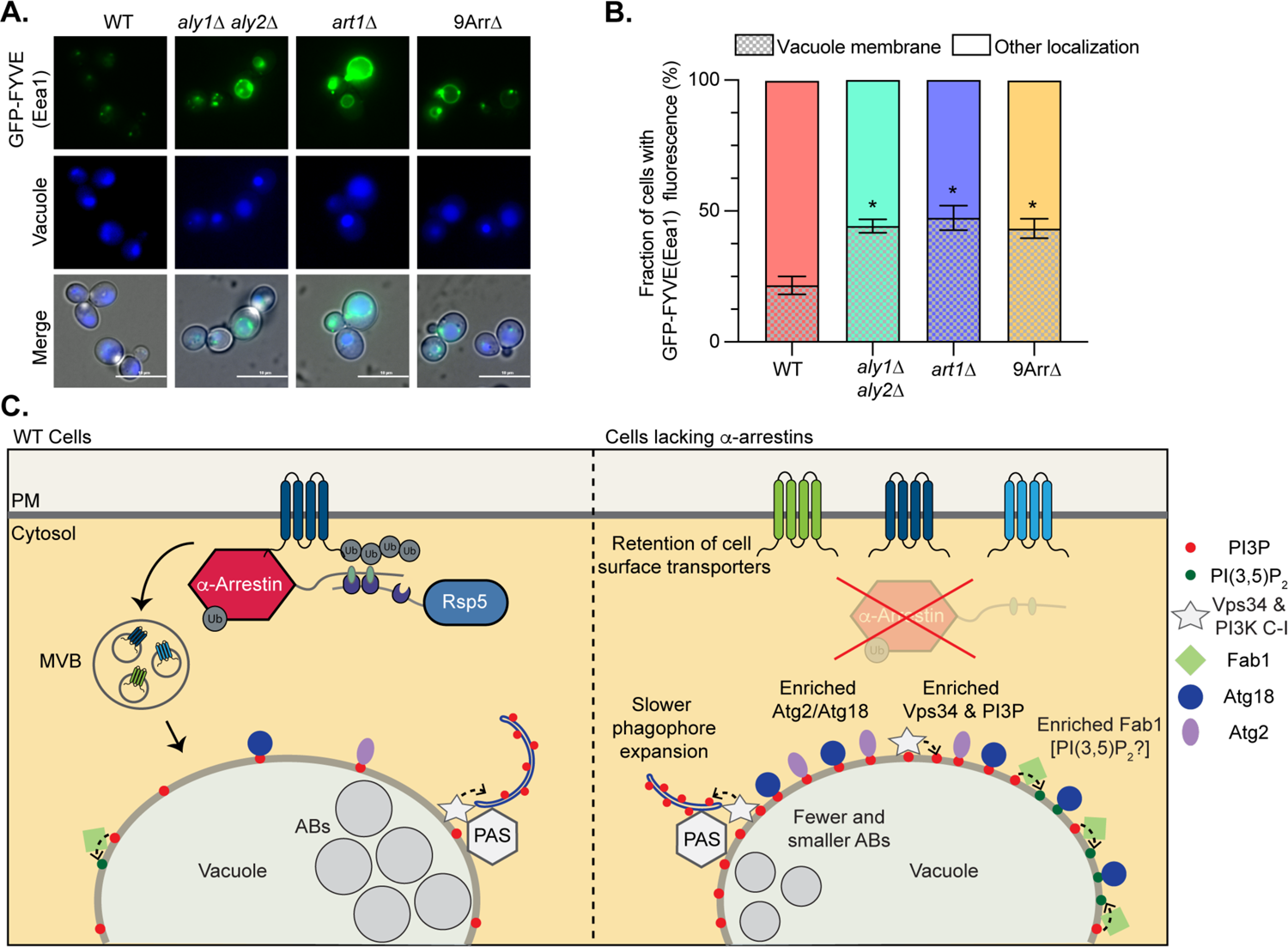
α-Arrestins are required for the proper distribution and abundance of PI3P. (A) Cells expressing pRS426-GFP-Eea1-FYVE [3] in either WT or α-arrestin mutant cells were imaged by fluorescence microscopy. The vacuole lumen was stained with CMAC. (B) The fraction of cells displaying GFP-Eea1-FYVE localization (as imaged in A) to the limiting membrane of the vacuole or to endosomes/other localizations was quantified for 4 replicate experiments using Image J. The median value is shown as a black line in the stacked bar graph and the error bars represent the 95% confidence interval. Unpaired Student’s t-test with Welch’s correction was used to assess statistical significance (* = p-value <0.05). (C) Model of α-arrestin’s influence on autophagy. When WT cells are treated with rapamycin, α-arrestins bind Rsp5 to control trafficking of membrane proteins to the vacuole. TORC1 signaling is impaired and autophagy is induced. Atg18-Atg2 bind to the vacuole membrane and are recruited to the PAS where AP expansion and subsequent fusion to the vacuole delivers autophagic bodies (AB) to the lumen of the vacuole for degradation. When cells lacking α-arrestins are treated with rapamycin, membrane transporters are aberrantly retained at the PM, resulting in loss of materials delivered to the vacuole. We find accumulation of PI3P and its kinase, Vps34, as well as Fab1, the kinase that generates PI(3,5)P_2_, at the vacuole membrane. Atg18 and its binding partner Atg2, each of which binds PIPs, is also enriched at the limiting membrane of the vacuole. This may interfere with their activity at the PAS, as we observe slower phagophore expansion and delivery of fewer and smaller Abs to the vacuole lumen.

## Discussion

Here we define the genetic interactions network between α-arrestins and the autophagy machinery. We find that while α-arrestin over-expression induces resistance to the TORC1 inhibitor, rapamycin, loss of key autophagy components altered this phenotype. Most strikingly, defects in components associated with the PAS, PI3K complex, and expansion of the phagophore membrane have the strongest genetic interactions with α-arrestins (Fig 2C) [45,47,50,51,73,75,136]. Our results agree well with the downstream disruptions to the autophagic machinery in cells lacking α-arrestins. Specifically, in cells lacking Aly1/Aly2, Art1 or 9 of the 14 α-arrestins (9ArrΔ), PI3P was more prevalent at the vacuole membrane (Fig 10A-C) [45, 137]. Vps34, the kinase that generates PI3P, was also enriched at the vacuole membrane in cell lacking α-arrestins. While there are no probes to directly examine PI(3,5)P_2_ localization in cells, we found increased Fab1, the PI3P kinase that generates PI(3,5)P_2_ [83,125,126], on the vacuole membrane when α-arrestins were absent, raising the possibility that there may also be more PI(3,5)P_2_ at this locale (Fig 10C). The key autophagy proteins that bind PI3P and/or PI(3,5)P_2_, Atg18 and Atg2, were also enriched on the limiting membrane of the vacuole when α-arrestin function was impaired. We posit that this is due to heightened accumulation of their phospholipid binding partners [60,61,83,122,124]. Consistent with a key role for PI3P enrichment in the delayed autophagic flux to the vacuole observed for cells lacking α-arrestins, we found that loss of α-arrestins prolonged AP lifespan, as has been observed in cells containing hyperactive Vps34 mutations [54]. Further, we find delayed expansion of the autophagosome membrane in cells lacking α-arrestins and reduced and/or smaller autophagic bodies accumulating in the vacuole of cells lacking these trafficking adaptors. Given the role for the Atg18-Atg2 complex in regulating lipid transfer at the PAS to help expand the phagophore [47,50,51], it is not surprising that this process is delayed in cells lacking α-arrestins, where aberrant Atg18-Atg2 enrichment along the entire vacuole membrane may prevent this complex from properly anchoring to the ER and/or concentrating as an active complex at the PAS.

Despite altered Atg18-Atg2 distribution, we saw no obvious defect in Atg9 levels or its recruitment to the PAS. Atg9 is a conserved transmembrane protein that arrives at the PAS via Atg9-containing vesicles, which have been proposed to be the ‘seed’ membrane source for phagophore formation [56,84,111,112]. Atg9-containing vesicles are formed at the Golgi and Atg27 and Atg23, a transmembrane and peripheral membrane protein, respectively, that complex with Atg9, are needed to generate these structures [113]. Atg27 trafficking is dependent upon adaptor protein complex 3 (AP-3), and in cells lacking AP-3, Atg9 accumulates in the vacuole lumen because of defective Atg27 trafficking [114]. We found that AP-3 mediated trafficking was unperturbed in cells lacking α-arrestins. Consistent with this idea, we did not observe accumulation of Atg9 in the vacuole lumen. Atg9 is integrated into the growing phagophore but is likely recycled from the PAS since it does not localize to the vacuole membrane. This recycling is thought to require PI3P, PI3K complex, Atg18-Atg2, and Snx4, a member of the SNX-BAR protein family, as well as retromer [84]. The Snx4 sorting nexin likely impacts autophagy through multiple modes; Snx4 in complex with Atg20 is thought to be important for endosome-to-Golgi trafficking of Atg9, which occurs via multiple routes [138, 139], but more recent work demonstrates that vacuole-to-endosome recycling of Atg27 relies specifically on Snx4-Snx41[132] and endosome-to-Golgi trafficking requires retromer [133]. While we did not assess Snx4 directly, we found that the residence of retromer and one of its clients, Vps10, were unchanged in the absence of α-arrestins. We did not observe any defects in Atg9 levels or recruitment to the PAS, but given the complexity of Atg9 and Atg27 recycling, the possible role for α-arrestins in protein recycling, and the recent findings that hyperactivated Vps34 increases Atg27 recycling [21,54,140–142], more nuanced studies are warranted in the future.

Given that loss of α-arrestins increased PI3P on the limiting membrane of vacuoles, recent studies on hyperactive Vps34 and its impact on downstream membrane trafficking and autophagy are relevant to our findings [54]. Vps34 forms the catalytic core of two distinct PI3K complexes and is the only kinase able to generate PI3P [78,137,143]. The first PI3K complex, known as Complex I, is required for autophagy and is comprised of Vps15, a pseudokinase and essential activator of Vps34, along with Atg6 (aka Vps30), Atg14 and Atg38 [79,144,145]. The second, known as Complex II, acts during many facets of protein trafficking and contains Vps34, Vps15, Atg6/Vps30 and Vps38 [45]. Though *VPS34* was not included in our genetic analyses, loss of *ATG6* resulted in rapamycin resistance. Interestingly, Aly over-expression in this background caused sensitivity, rather than resistance, to this TORC1 inhibitor (Fig 2B and S2 Fig). In cells lacking *ATG6*, the catalytic function of both PI3K complexes is impaired, leading to reduced PI3P in cells and impeding autophagic flux [78, 146]. This result suggests that whatever positive influence Aly over-expression has on TORC1 function, this effect is reversed in the absence of *ATG6*.

Unlike the strong genetic interaction with *ATG6*, loss of the autophagy specific component of Complex I, *ATG14*, had a more modest impact on Aly-mediated resistance to rapamycin. Cells lacking *ATG14* were resistant to rapamycin and over-expression of Alys further increased this resistance, indicative of a positive genetic interaction. These data suggest that Alys’ influence on rapamycin resistance may be more dependent on Complex II activities than the autophagy specific Complex I.

Vps34 and PI3P are needed for retromer-dependent retrieval of membrane cargo from late endosomes to the early endosome, generation of PI(3,5)P_2_, ESCRT function and MVB fusion to the vacuole, along with Snx4-mediated retrieval of membrane proteins from the vacuolar membrane [54,58,59,125,126,133,134]. As mentioned above, hyperactive Vps34 increases either Snx4- and/or retromer-dependent recycling of Atg27 from vacuoles [54], yet retromer function was unaltered in cells lacking α-arrestins. Thus, while some elements are consistent between Vps34 hyperactivation and the phenotypes associated with loss of α-arrestins, they are imperfectly aligned. Deletion of the genes encoding α-arrestins delays endocytic turnover of many plasma membrane transporters [15,16,18,22,34], and ESCRTs operate synergistically downstream of the α-arrestins to help ensure these membrane proteins reach the vacuole lumen. Thus, to address ESCRT function in the absence of α-arrestins, one would need an α-arrestin-independent membrane cargo as a readout, but these studies could still be confounded by the severe disruption to protein trafficking that is caused by α-arrestins.

Hyperactive Vps34 increases PI(3,5)P_2_ production, which is particularly apparent in response to osmotic shock [54]. While there is no probe to measure PI(3,5)P_2_ levels, we did find that Fab1 accumulates on the vacuole membrane and this raises the possibility that there is elevated Fab1 activity in the absence of α-arrestins. This elevated Fab1 at the vacuole is different from what was observed for the hyperactive Vps34 mutant, which did not appear to accumulate increased Fab1 [54]. Finally, hyperactive Vps34 mutations increased GFP-Atg8 puncta lifetime and reduced GFP-Atg8 breakdown in the vacuole post nitrogen starvation [54]. Similarly, in our rapamycin treated cells, loss of α-arrestins increased GFP-Atg8 puncta lifetime, delayed GFP-Atg8 delivery to the vacuole and reduced autophagic body accumulation in the vacuole. However, when Atg18 localization was examined in hyperactive Vps34 cells, there was no increase in Atg18 accumulation on vacuole membranes but there was a defect in the disassembly of Atg18 from APs [54]. Thus, hyperactive Vps34 and elevated PI3P delayed AP fusion/completion and this is thought to be due, in part, to a more general defect in homotypic fusion of vacuoles, as supported by an increase in the number of vacuole lobes in Vps34 hyperactive cells [54]. Like Vps34 hyperactivated cells, *aly1*Δ *aly2*Δ cells have hyper-fragmented vacuoles. However, we observe enlarged vacuoles in the *art1*Δ or 9ArrΔ cells, a phenotype associated with loss of Atg18 or defects in PI(3,5)P_2_ levels, and so there must be additional drivers of the vacuole morphology in these strains other than simply elevating PI3P abundance. The enlarged, single lobed vacuoles of *art1*Δ and 9ArrΔ are consistent with possible elevated vacuole fusion or impaired vacuole fission, which remains to be further explored in subsequent studies.

Countering Vps34 activity are the <u>y</u>east <u>m</u>yotubulinar-related PI3P phosphatase, Ymr1, and the <u>s</u>ynaptojanin-like phosphatases, Sjl2 and Sjl3 [147], which may help return PI3P back to PI [148, 149]. Loss of Ymr1 and Sjl2 and Sjl3 is synthetically lethal, but this defect can be rescued by targeting the catalytic core of Sac1, a PIP phosphatase, to PI3P-containing membranes by creating a chimeric protein with the Eea1-FYVE domain. This suggests that the cause of *ymr1*Δ *sjl2*Δ *sjl3*Δ lethality is excess accumulation of PI3P [147]. Indeed, cells defective for Ymr1 and Sjl2 activity have >2-fold higher PI3P levels, and, as observed with the α-arrestin mutants, this PI3P aberrantly accumulates on the vacuolar membrane. Interestingly, *ymr1*Δ *sjl2*Δ cells, in marked contrast to the *art1*Δ or 9ArrΔ cells, have hyper-fragmented vacuoles that is more reminiscent of hyperactive Vps34 or *aly1*Δ *aly2*Δ cells [147]. Thus, while each of these mutants appears to accumulate more PI3P on the vacuole membrane, based on increased localization of the fluorescent FYVE domain of Eea1, there must be other distinguishing features of these mutants that give rise to these opposing vacuole morphologies. Perhaps these altered morphologies represent varying levels of PI3P at the vacuole or, more likely, they are due to multifactorial inputs. In addition to these disparate vacuole morphology changes, cells defective for Ymr1 and/or Sjl2 function have impaired retromer function, fail to fuse APs to the vacuole membrane and fail to dissociate Atg18 from APs [147, 150]. We did not observe any obvious defects in retromer function (i.e., Vps10 sorting was intact and Vps17 localization appeared normal), and in cells hyperactivated for Vps34 there was similarly no defect in retromer function [54]. Because we see dramatic changes in Atg18 localization to the limiting membrane of the vacuole, we suspect this is linked to defects in AP biogenesis.

While Atg18 plays a key role in AP formation, it also regulates vacuole morphology by interacting with the PI3P kinase, Fab1, and its regulator Vac14, which also associate with retromer to control vacuole fragmentation [46,52,83,151]. Cells lacking Atg18 have enlarged vacuoles with excessive PI(3,5)P_2_, which is somewhat surprising since vacuoles depleted for PI(3,5)P_2_ (as happens in Fab1 hypomorphic mutant cells) also have enlarged, unilobed vacuoles [152, 153]. Perturbations in the PIP balance at the vacuole membrane drives many vacuole morphology defects, likely due to changes in the regulation of the vacuole fission and fusion machinery. For example, in response to hyperosmotic shock, vacuoles fragment and this process requires PI3P and the action of Vps34 followed by the action of PI(3,5)P_2_, Fab1 and Atg18 [61], demonstrating the complex interplay that occurs between PIPs and their regulators to control vacuole morphology. At the vacuole, Atg18 recruitment requires Fab1 and Vac7 or Vac14, each of which are Fab1 regulators [83]. We find that Fab1 is enriched on the vacuole membrane in the absence of select α-arrestins, which could aid in Atg18 recruitment to this site. Enlarged vacuole morphologies, like those displayed by *art1*Δ or 9ArrΔ cells, are reminiscent of the Class D vacuoles first described by the Steven’s lab [154]. Class D mutant vacuoles often contain markers for both endosomes and the vacuole and have impaired vacuole fission [155, 156]. Indeed, defects in vacuole morphology often impede vacuole inheritance, yet another vacuole-based process for which Atg18 is needed [83, 157].

Most recently, the identification of Vps35 in proximity proteomics studies of Atg18 highlights a new facet of Atg18’s regulation of vacuole morphology [52]. Atg18 co-purifies with Vps35, and this association appears to be competitive with Vps35 binding to the sorting nexin pair, Vps5-Vps17, which are normally associated with its function in recycling proteins from endosomes to the Golgi along with Vps26, an arrestin-like protein, as part of retromer [52,158,159]. These findings suggest that Atg18 may provide the membrane binding capacity for a new variant of retromer, where Atg18 replaces Vps5-Vps17 [52]. The Atg18-Vps35 complex is needed for vacuole fragmentation in hyperosmotic stress, shedding new light on the vacuole fission process [52]. Cells lacking Vps35 have enlarged vacuoles and increased Atg18 localization to the limiting membrane [52], which are similar to the phenotypes we see in *art1*Δ and 9ArrΔ cells. Given that retromer contains an arrestin-like protein, Vps26, it is tempting to speculate about the ability of other α-arrestins to substitute for Vps26 in this new variant of the retromer complex. It will be exciting to define the role of α-arrestins in vacuole inheritance and/or regulation of vacuole fission in future work as it seems likely, given the abnormal vacuole morphologies and altered distributions of Atg18, Fab1 and other regulators, that α-arrestins will play an important role in for these activities.

Ultimately, it will be critical to determine how the loss of α-arrestins alters PIP distribution. PIP distributions could change because cellular metabolite balance is altered when α-arrestins are missing. In the absence of α-arrestins, nutrient transporters are retained at the cell surface and this likely disrupts cellular levels of key metabolites, which might in turn affect PIP biosynthesis. For example, Aly1 and Aly2 regulate the trafficking of the glycerophosphoinositol (GPI) transporter, Git1, that allows GPI into cells [160]. In the absence of Alys, Git1 accumulates at the cell surface and cells become exquisitely sensitive to GPI overload, suggesting that excess GPI is toxic [160]. Once inside cells, GPI is converted into glycerol-3-phosphate and inositol [161, 162], which is a building block for PIP synthesis. Indeed, our earlier studies showed that vacuole membranes in cells lacking Alys have elevated PI3P, and the over-abundance of PI3P is exacerbated by addition of exogenous GPI [160]. In another example, inositol uptake is controlled by Itr1 and Itr2, and α-arrestin Art5 recognizes Itr1, and perhaps Itr2, to regulator inositol-induced endocytosis [16, 163]. In the absence of Art5, cellular inositol levels likely rise due to Itr retention at the PM and unregulated inositol uptake. In fact, mis-regulation of Itr1 endocytosis alters mitochondrial cardiolipin production and can rectify defects in phospholipid flippases [163–165]. Interestingly, like Aly1 and Aly2, Art5 over-expression conferred rapamycin resistance to cells (S2A Fig). These connections raise the possibility that the rapamycin resistance and regulation of inositol homeostasis may be linked to the changes in PI3P and its binding partners at the vacuole membrane.

In addition to a possible connection between inositol balance, PIPs and α-arrestin-mediated trafficking, other metabolite perturbations are linked to enlarged vacuoles and Atg18 sequestration on the membrane: treatment of cells with methylglyoxal, a metabolic intermediate in a glycolytic bypass pathway, leads to cellular toxicity, increases vacuole size, retains of Atg18 on the limiting membrane and prevents hyperosomotic-induced fragmentation of vacuoles [166, 167]. Given the broad roles of α-arrestins in controlling membrane protein trafficking, their impact on cellular metabolism is likely also far reaching and complex. Future studies aimed at defining the metabolic impact of α-arrestin mutants would undoubtedly help us in defining links between regulated protein trafficking, phospholipid balance and maintenance of organelle morphology/function.

Finally, in the model presented above (Fig 10), the role of α-arrestins in regulating vacuole morphology and composition could be linked to the protein trafficking of plasma membrane proteins and subsequent nutrient balance changes associated with defects in that process. However, it is formally possible that the α-arrestins may directly regulate the vacuole membrane proteome and its composition analogous to what has been shown for other Rsp5 ubiquitin ligase adaptors, including Ssh4 or Ear1 [168–171]. In sum, our work highlights important areas of future study that are likely to further expand our appreciation of how regulated protein trafficking impacts metabolism and organelle function.

## Materials and Methods

### Yeast strains and growth conditions

The yeast strains used in this study are described in S1 Table and are derived from the BY4741 genetic background of *S. cerevisiae* (S288C in origin). Yeast cells were grown in either <u>s</u>ynthetic <u>c</u>omplete (SC) medium lacking the appropriate amino acid for plasmid selection and using ammonium sulfate as a nitrogen source [172] or <u>y</u>east extract <u>p</u>eptone <u>d</u>extrose (YPD) medium where indicated. Unless otherwise indicated, yeast cells were grown at 30°C. Liquid medium was filter-sterilized and solid medium for agar plates had 2% agar w/v added before autoclaving.

### Plasmids and DNA manipulations

Plasmids used in this work are described in S2 Table. PCR amplifications for building plasmid DNA inserts were performed using Phusion High Fidelity DNA polymerase (ThermoFisher Scientific, Waltham, MA) and confirmed by DNA sequencing. Plasmids were transformed into yeast cells using the lithium acetate method [173] and transformants were selected for on SC medium lacking a specific nutrient.

### Yeast protein extraction and immunoblot analyses

Whole-cell extracts of yeast proteins were prepared using the tri<u>c</u>holoro<u>a</u>cetic acid (TCA) extraction method as previously described [71] and as modified from [174]. In brief, cells were grown in SC medium to mid-exponential log phase at 30°C (A_600_= 0.6-1.0) and an equal density of cells was harvested by centrifugation. Cell pellets were flash frozen in liquid nitrogen and stored at −80°C until processing. Cells were lysed using sodium hydroxide and proteins were precipitated using 50% TCA. Precipitated proteins were solubilized in SDS/Urea sample buffer [8M Urea, 200mM Tris-HCl (pH 6.8), 0.1mM EDTA (pH 8), 100mM DTT, Tris 100mM (not pH adjusted)] and heated to 37°C for 15 minutes. Samples were then precipitated using 50% TCA and solubilized in SDS/Urea sample buffer as above. Proteins were resolved by SDS-PAGE and identified by immunoblotting with a mouse anti-<u>g</u>reen fluorescent <u>p</u>rotein (GFP) (Santa Cruz Biotechnology, Santa Cruz, CA), or anti-Atg13 [175] antibody to detect tagged or endogenous proteins. As a protein loading control, immunoblot membranes were stained after transfer with Revert^TM^ (Li-Cor Biosciences, Lincoln, NE) total protein stain and detected using the Odyssey^TM^ CLx Infrared imaging system (Li-Cor Biosciences). Anti-mouse or anti-rabbit secondary antibodies, conjugated to IRDye-800 or IRDye-680 (Li-Cor Biosciences) were used to detect primary antibodies on the Odyssey^TM^ CLx (Li-Cor Biosciences).

### Serial dilution growth assays

For the serial dilution growth assays on solid medium, cells were grown to saturation in liquid YPD or SC medium overnight and the A_600_ determined. Starting with an A_600_ of 1.0 (approximately 1.0 x 10^7^ cells/ml), 5-fold serial dilutions were generated and transferred to solid medium using a sterile replica-pinning tool. Cells were grown at 30°C for 3-6 days and images captured using a Chemidoc XRS+ imager (BioRad, Hercules, CA). All images were evenly adjusted in Photoshop (Adobe Systems Incorporated, San Jose, CA). For rapamycin (LC Laboratories, Woburn, MA) containing plates, the stock solution of 0.5 mg/ml in ethanol was diluted in growth medium to the final concentration indicated in each figure panel (typically 50 ng/ml).

### ScUbI library screen

The *<u>S</u>accharomyces <u>c</u>erevisiae* <u>Ub</u>iquitin Interactome (ScUbI) library contains 323 unique non-essential gene deletions, each of which is annotated as being non-essential in the *<u>S</u>accharomyces* <u>G</u>enome <u>D</u>atabase (SGD). This library initially was constructed as the <u>T</u>argeted <u>U</u>biquitin <u>S</u>ystem (TUS) yeast gene deletion library [176] We then used YeastMine, populated by SGD and powered by InterMine, and searched for ‘ubiquitin’. These searches returned over 2,000 candidates before duplicates, essential genes, and targets of ubiquitin were removed. The remaining list was combined with the TUS library to produce the final ScUbI library. The 323 gene deletion mutants were arrayed over four 96-well plates. The library is available upon request and is further documented at https://www.odonnelllab.com/yeast-libraries.

To screen for modifiers of α-arrestins Aly1 and Aly2 function, the ScUbI library was transformed with either pRS426-vector or pRS426-Aly1 or -Aly2 plasmids, each of which over-express Aly1 or Aly2 as they were present on a 2-micron plasmid backbone, but expressed from their endogenous promoter. Transformations were performed as described in [173] and were done with the aid of the Benchtop RoToR HAD robotic plate handler (Singer Instruments Co. Ltd, Roadwater, UK) and the Multidrop Combi (ThermoFisher Scientific, Waltham, MA) in the laboratory of Dr. Anne-Ruxandra Carvunis (Univ. of Pittsburgh). Each transformed version of the library was stamped in technical triplicate to SC medium lacking uracil (as a control), or the same medium containing 50 ng/ml rapamycin. Plates were grown at 30°C for 2-4 days and white light images were captured using the BioRad ChemiDoc XRS+ imager (Hercules, CA) on days 1-4 of incubation. Images were converted to .*jpg* file format and the pixel size of colonies were measured using the DissectionReader macro [generously provided by Dr. Kara Bernstein’s laboratory (Univ. of Pittsburgh) and developed by John C. Dittmar and Robert J.D. Reid in Dr. Rodney Rothestein’s laboratory (Columbia Univ.)] in Image J (National Institutes of Health, Bethesda, MA, USA). More information on this plugin can be found at: https://github.com/RothsteinLabCUMC/dissectionReader. Colony sizes were converted into individual sets of Z-scores for each transformed version of the library (this allowed for comparisons between pRS426 vs pRS426-Aly1 or -Aly2 colony sizes). The Z-scores for each gene deletion strain containing pRS426 was subtracted from the Z-score for that gene deletion when over-expressing Aly1 or Aly2 to produce the ‘change from vector’ or ΔV value (S1 Datafile). A ΔV cutoff of +/- 1.0 was arbitrarily used to identify a targeted list of gene deletion candidates for further study. This list was used to perform <u>G</u>ene <u>O</u>notology (GO) enrichment analysis described below.

### Gene ontology enrichment analysis

GO trees (file: *go-basic.obo*) and annotations (files: *sgd.gaf*) were downloaded from http://geneontology.org/ on April 20, 2022. We used *BINGO* from the *Cytoscape* application library to calculate the number of genes associated with each GO term in the group with strongest Z-score change (positive or negative) and the overall population of (all) genes we tested (see *ScUbI Library Screen* above, S1 Datafile) [177, 178]. We excluded annotations based on the evidence codes ND (no biological data available). We identified GO term enrichments by calculating the likelihood of the ratio of the genes associated with a GO term within a study group given the total number of genes associated with the same GO term in the background set of all genes in our population. We applied a hypergeometric test to calculate *p*-values for the enrichment of GO terms. *p*-value < 0.05 was taken as a requirement for significance.

To visualize GO terms, we removed redundant information by selecting parent GO terms of multiple similar terms. Then we calculated the average Z-score change for the genes associated with the GO term and plotted representative GO terms based on the associated p-values and average Z-score changes using R package ggplot2 v3.3.5 [179].

### Quantification of serial dilution growth assays

Serial dilution growth assays in S1-2 Figs) were quantified using a CellProfiler (Broad Institute, Cambridge, MA) automated pipeline (available at https://www.odonnelllab.com/automated-quantification-pipelines). White light images of serial dilution growth assay plates were obtained using a BioRad ChemiDoc XRS+ imager (Hercules, CA, USA) after 5 days growth. Images were then cropped, converted to binary, and resized to equivalent pixel densities. The CellProfiler automated pipeline segmented the rows and columns for each image and measured the pixel intensity for each spot of yeast growth, aggregating the values for each row. Using this approach, we defined a quantitative measure of the growth present for each row in a serial dilution growth assay. These values were converted into a heat map using Prism (GraphPad Software, San Diego, CA). Data derived from this analysis is presented in Fig 2B and S2C Fig.

### Electron Microscopy

We grew cells overnight in SC medium in culture tubes on a rotating drum at 30°C. We then diluted cells (∼A_600_ = 0.3) and regrew them in culture tubes for 4 h on a rotating drum at 30°C. For cells treated with rapamycin (LC Laboratories, Woburn, MA), 200 ng/ml rapamycin was added to 15ml of cells, then incubated at 30°C for four hours. We used an osmium tetroxide (OsO4) substitution method to prepare yeast cells for EM [180]. Concentrated live yeast cells were placed in 1.5 mm wide by 0.2 mm deep brass planchets (Leica 707899) and frozen using a Leica EM PACT2 High pressure freezer (Leica Microsystems, Buffalo Grove, IL). Frozen samples were then placed in a Leica EM AFS2 Freeze Substitution device (Leica Microsystems, Buffalo Grove, IL). A solution of 2% OsO_4_ in 100% acetone was placed on the samples for 110 hours at minus 90°C. The samples were then left in the OsO_4_ solution, and the temperature was allowed to rise 5°C every hour until the solution reached 0°C. Samples remained at 0°C and were dehydrated further with 3 x 30-minute acetone washes and 3 x 1-hour ethanol washes (Pharmco, Brookefield, CT). Samples were then removed from the Freeze Substitution device and allowed to come to room temperature. Samples were washed twice in propylene oxide (Electron Microscopy Sciences, Hatfield, PA) and embedded in Poly/Bed® 812 (Luft formulation, Warrington, PA). Samples were allowed to cure for 24 hours at 37°C and 48 hours at 65°C. The brass planchets were carefully removed from the polymerized resin and semi-thin (300 nm) sections were cut on a Leica Reichart Ultracut (Leica Microsystems, Buffalo Grove, IL), stained with 0.5% Toluidine Blue in 1% aqueous sodium borate (Fisher, Pittsburgh, PA) and examined using light microscopy. Ultrathin sections (65 nm) were taken, placed on 200 mesh copper TEM grids and were stained with 2% uranyl acetate (Electron Microscopy Sciences, Hatfield, PA) and Reynold’s lead citrate (Fisher, Pittsburgh, PA) and examined on JEOL 1400 Plus transmission electron microscope (JEOL Peabody, MA) with a side mount AMT 2k digital camera (Advanced Microscopy Techniques, Danvers, MA).

### Quantification of Electron Micrographs

The number of autophagic bodies (AB) per cell was determined manually by counting the number of membrane-bound structures inside the vacuoles of complete cells for each electron micrograph. The diameter of ABs measurement was performed using Image J software (NIH, Bethesda, MD). The distance scale was calibrated using the scale bar on the electron micrograph and the furthest distance across each AB was determined. Data derived using this method is presented in Fig 3B-C.

### Fluorescence microscopy

Fluorescent protein localization was assessed using epifluorescent and confocal microscopy. We grew cells overnight in SC medium in culture tubes on a rotating drum at 30°C. We then diluted cells (∼A_600_ = 0.3) and regrew them in culture tubes for 4 h on a rotating drum at 30°C. For cells treated with rapamycin (LC Laboratories, Woburn, MA), 200 ng/ml rapamycin was added to 1ml of cells in a microcentrifuge tube and incubated in an end-over-end rotator at 30°C for indicated times. Prior to imaging, FM4-64 or CMAC stains were used to mark the vacuole membrane or lumen, respectively. For cells stained with FM4-64 (ThermoFisher Scientific, Waltham, MA), 1 ml of cells were treated with 100 μg/ml FM4-64 for 30 minutes at 30°C. Cells were then washed, resuspended in 1 ml SC medium, and incubated at 30°C for 1h. For cells stained with CMAC, 20 μM Cell Tracker Blue CMAC (7-amino-4-chloromethylcoumarin) dye (Life Technologies, Carlsbad, CA) was added to 1 ml of mid-log cells 1 hour prior to imaging. Cells were then inoculated to low density (∼A_600_ = 0.15) onto 35 mm glass bottom microwell dishes (MatTek Corporation, Ashland, MA) that were either poly-D-lysine coated or had been treated with 50 μl of 0.2 mg/ml concanavalin A (MP Biomedicals, Solon, OH). Cells were imaged by confocal microscopy using a Nikon Eclipse Ti2-E A1R inverted microscope (Nikon, Chiyoda, Tokyo, Japan) outfitted with a 100x objective (NA 1.49) and images were detected using GaAsP or multi-alkali photomultiplier tube detectors as single median planes or as a series of images in an 8-10 μm Z-stack with 0.125-0.25 μm step sizes. Data derived using this method is presented in Figs 4-9, and S4,7-8 Figs. Alternatively, cells were imaged by epifluorescent microscopy using a Nikon Eclipse Ti2 inverted microscope (Nikon, Chiyoda, Tokyo, Japan) outfitted with a Teledyne Photometics Prime BSI CMOS (sCMOS) camera (Teledyne Photometics, Tucson, AZ) and a 100x objective (NA 1.45). Data derived using this method is presented in Fig 10A, and S5 Fig. In both cases, acquisition was controlled using NIS-Elements software (Nikon, Chiyoda, Tokyo, Japan) and all images within an experiment were captured using identical settings. Confocal images were deconvolved using the Richardson-Lucy algorithm. All images were cropped and adjusted evenly using NIS-Elements.

### Fluorescent image quantification and statistical analyses

Quantification of whole cell fluorescence intensity was done using the Nikon General Analysis 3 software (Nikon, Chiyoda, Tokyo, Japan) using segmentation from NIS-Elements.*ai* (Artificial Intelligence) software (Nikon, Chiyoda, Tokyo, Japan) unless otherwise described below. For quantification of whole-cell signal, the NIS.*ai* software was trained on a ground truth set of samples where cells had been manually segmented using DIC channel images. Next, the NIS.*ai* software performed iterative training until it achieved a training loss threshold of <0.02, which is indicative of a high degree of agreement between the initial ground truth provided and the output generated by the NIS.*ai* software. Fields of images captured via confocal imaging were then processed so that the individual cells in a field of view were segmented using the DIC. Any partial cells at the edges of the image were removed. The sum fluorescence intensity and pixel count for each cell was defined in the appropriate channel, with the mean fluorescence intensity being calculated by dividing the sum intensity by the number of pixels. Data derived from these kinds of analyses are presented in Fig 5 and S4 Fig.

To measure the vacuolar lumen or membrane fluorescence or length, we trained the NIS.*ai* software to identify the vacuolar lumen or membrane using CMAC or FM4-64, respectively, as fiducial markers. The NIS.*ai* software was trained using a manually defined ‘ground truth’ set of vacuole segmentations selecting either the entire vacuole or just the vacuolar membrane and trained as above. Then, fields of images captured via confocal imaging were processed so that the individual vacuole objects in a field of view were segmented using the 405nm (CMAC) or 561nm (FM4-64) channel. Using General Analysis 3 software, individual cells were segmented by the same trained NIS.*ai* module described above using the DIC channel. A parent-child relationship was applied to individual vacuolar objects (child) within the same cell (parent) to aggregate them as single objects and pair them to the appropriate whole cell. Any partial cells at the edges of the image were removed along with their child-objects. Then the sum fluorescence intensity and pixel count for each parent or child object was defined in the appropriate channel. Mean intensities were determined by dividing the sum intensity by the number of pixels for all child objects inside a parent. Mean vacuolar lengths were determined by dividing the sum intensity by the sum pixel counts for each child object inside a parent. Data derived from these kinds of analyses are presented in Figs 7-9, and S6-8 Figs.

Manual image quantification to measure the vacuolar membrane fluorescence or length was performed using ImageJ software (National Institutes of Health, Bethesda, MD). A 2-pixel thick line was hand drawn over the vacuolar membrane using images captured of cells stained with FM4-64 which was then overlaid on the GFP images and the median GFP signal or total length of the line was measured. The median background fluorescence intensity was then subtracted for the vacuolar membrane intensity measures. Data derived from these kinds of analyses are presented in Fig 7 and S6 and S8 Fig. Manual image quantification to measure the number of vacuole lobes was performed using ImageJ software (National Institutes of Health, Bethesda, MD). The number of vacuole lobes was determined using images captured of cells stained with FM4-64. Data derived from this kind of analysis is presented in S6C Fig. To determine the lifetimes of GFP-Atg8 AP structures, we used time-lapse confocal microscopy to obtain Z-stack image series acquired every 30 seconds for 20-25 minutes per single field of cells. Following Richardson-Lucy deconvolution, images were manually assessed to identify newly formed GFP-Atg8 puncta which were then tracked over time until they merged with the vacuole (stained with CMAC), recording the total lifetime of the puncta. Representative cells displayed as maximum intensity projections of the entire Z-series (Fig 4), or as 3D projection movies (S1-4 Movies) were created using NIS Elements software.

To measure Atg9-mNG intensity at the PAS, we relied on the autophagy cargo protein Ape1, which is recruited to the PAS upon autophagy induction [57]. We trained the NIS.*ai* software to segment the PAS using the signal from puncta formed by mCherry-Ape1 expressing cells. The NIS.*ai* software was trained using a manually defined ‘ground truth’ set of mCherry-Ape1 PAS segmentations as above. Fields from images captured via confocal imaging were then processed so that individual PAS objects in a field of view were segmented using the 561nm channel. Parent-child relationships were used to pair PAS to their respective whole as above, and any partial cells at the edges of the image were removed along with their child-objects. The sum fluorescence intensity for each PAS was defined in the 488nm channel to assess the abundance of Atg9-mNG recruited following 3h rapamycin treatment. Data derived from this analysis is presented in Fig 6D.

To evaluate the ability of isolation membranes to expand, we utilized the giant Ape1 assay [88, 181]. We used confocal microscopy to obtain Z-stack image series acquired following 1h treatment with rapamycin. Following Richardson-Lucy deconvolution, BFP-Ape1 puncta were manually assessed for the presence of GFP-Atg8 structures. When present, these structures were manually binned into categories of ‘patch’ and ‘elongated’ based on their shape and size. Data derived from this analysis is presented in Fig 6E.

Fluorescent quantification was assessed statistically using Prism (GraphPad Software, San Diego, CA). Unless otherwise indicated, we performed the Kruskal-Wallis statistical test with Dunn’s post hoc correction for multiple comparisons. In all cases, significant p-values from these tests are represented as: *, p value < 0.1; ** p value < 0.01; ***, p value < 0.001; ****, p value < 0.0001; ns, p value > 0.1. In some instances where multiple comparisons are made, the † symbol may additionally be used in place of the * with the same p value meanings but indicating comparisons to a different reference sample.

## Acknowledgments

This work was supported by National Science Foundation CAREER grant MCB 155143 and 1902859 to A.F.O., developmental funds from the Dept. of Biological Sciences (Univ. of Pittsburgh) to A.F.O., and the Norman H. Horowitz and Samuel D. Colella Undergraduate Summer Research Awards to E.M.J. Electron microscopy work was supported by NIH 1S10RR019003-01, 1S10RR025488-01, and 1S10RR016236-01 grants to Simon Watkins, the Director of the Center for Biologic Imaging. We gratefully acknowledge the work of the Introduction to Molecular Genetics and Advanced Cell and Molecular Biology laboratory classes at the Univ. of Pittsburgh and Duquesne University, respectively. We thank Lois Weisman (Dept. of Cell & Developmental Biology, Univ. of Michigan), Daniel Klionsky (Dept. of Molecular, Cell and Developmental Biology, Univ. of Michigan), Claudine Kraft (Dept. of Biochemistry II, Albert-Ludwigs Univ. of Freiburg), Nava Segev (Dept. of Biochemistry and Molecular Genetics, Univ. of Illinois Chicago), Marijn Ford (Dept of Cell Biology, Univ. of Pittsburgh), and Jeffrey Brodsky (Dept. of Biological Sciences, Univ. of Pittsburgh) for generously providing reagents to aid in these studies and useful scientific discussions. We thank Anne-Ruxandra Carvunis and Nelson Castilho Coelho for their assistance with the robotics needed for our ScUbI screen. We thank Christian Ungermann (Biochemistry section, Osnabruck Univ.) for helpful suggestions during these studies. We further thank members of the O’Donnell and Brodsky labs for their input during these experiments.

